# Conformational dynamics at the pre-miR-377 Dicer site governs selective small-molecule recognition

**DOI:** 10.64898/2026.02.11.705234

**Authors:** Jacopo Manigrasso, Emeline Mestdach, Riccardo Quaia, Daniela Dankova, Mattias Bood, Malin Lemurell, Stefan Geschwindner, Andrey I. Frolov, Matthew D. Disney, Anders Hogner, Celestine N. Chi, Giuseppina La Sala

## Abstract

Small molecules that target precursor microRNAs (pre-miRNAs) offer a strategy to modulate microRNA biogenesis, but rational design is limited by the intrinsic conformational dynamics of RNA. Here, we combine site-specific nuclear magnetic resonance (NMR) spectroscopy with enhanced-sampling molecular dynamics (MD) simulations to define the conformational ensemble of pre-miR-377 and its recognition by the small-molecule binder C1. Our data reveals that the Dicer cleavage site contains a dynamic adenosine bulge in which U25 shuffles between A3 and A4 with a ∼9:1 population ratio, generating two interconverting RNA triple structures. NMR-informed MD and binding simulations show that C1 engages this dynamic region through a pathway-dependent mechanism that favors minor-groove entry, stabilizes intermediate, partially reorganized Dicer-site conformations, and redistributes the conformational ensemble away from processing-competent states. A conformationally locked mutant abolishes binding, indicating that recognition depends on bulge-enabled RNA dynamics rather than a single static structure. These results define the structural and dynamic basis of small-molecule recognition at the pre-miR-377 Dicer site and highlight the importance of RNA conformational ensembles in RNA-targeted drug discovery.

## INTRODUCTION

MicroRNAs (miRNAs) are post-transcriptional regulators of gene expression and are involved in numerous cellular processes including development, differentiation, and apoptosis.^1–3^ As important figures in the gene regulatory network, miRNAs dysregulation is frequently associated with pathological conditions including cancer and cardiovascular disorders.^4,5^ The maturation of these small ∼22-nucleotide-long RNA molecules involves a highly controlled biogenesis pathway, encompassing two RNA processing events: the first is the nuclear cleavage of primary transcripts (pri-miRNAs) by the microprocessor complex to produce miRNA precursors (pre-miRNAs).^6^ Then, following nuclear export, pre-miRNAs are further cleaved by the Dicer endonuclease to generate mature miRNA duplexes that guide the RNA-induced silencing complex (RISC) in targeting specific messenger RNAs (mRNAs).

The multi-step nature of this pathway presents attractive opportunities for therapeutic intervention through small molecule targeting. By binding to specific RNA structures; such as pre-miRNA substrates that serve as Dicer recognition sites; small molecules can disrupt miRNA maturation and modulate downstream gene expression networks.^7–9^ This therapeutic strategy has demonstrated success across multiple disease-relevant targets, including the precursors of pathogenic miRNAs such as miR-96, miR-515, miR-885, miR-372, and miR-21.^10–16^ Together with the clinical success of Risdiplam; a small molecule splicing modulator used to treat spinal muscular atrophy; these advances exemplifies how small molecules can engage human RNAs with high specificity and therapeutic efficacy, validating RNA-targeted small molecule drug discovery as a viable and impactful strategy.^17,18^

The molecular determinants of RNA-small molecule recognition, however, remain poorly understood, in part due to the conformational plasticity inherent to pre-miRNAs. Indeed, such structural flexibility poses significant challenges for the atomic structural characterization of RNA-small molecule binding mechanisms,^19–21^ hampering conventional structure-based design of binders. Unlike proteins, which often present stable binding sites, RNA molecules often exist as dynamic ensembles of interconverting conformations that undergo functionally critical rearrangements across multiple timescales,^21–29^ challenging both conventional experimental structure determination and computational modeling approaches.^19,30–35^ As such, most RNA-targeting drug discovery campaigns rely on combinatorial screening or sequence-based approaches rather than 3D structure-based design principles.^36,37^ This knowledge gap is underscored by the scarcity of pre-mRNA-small molecule complexes in the Protein Data Bank,^38^ which limits our atomic understanding of pre-miRNA-small molecule interactions.

Here, efforts are focused on characterizing the recognition and binding mechanisms of small molecules targeting pre-miR-377, a promising therapeutic candidate for the treatment of cardiovascular diseases. The miR-377 is aberrantly upregulated in ischemic myocardium, where it pathologically represses expression of vascular endothelial growth factor A (VEGFA), thereby impairing angiogenesis and vascular repair.^39^ Recently, two-dimensional combinatorial screening (2DCS) coupled with Inforna sequence-based analysis identified a phenylbenzimidazole derivative, dubbed C1, that selectively binds the Dicer processing site of pre-miR-377 (hereafter referred to as Dicer site) and inhibits its processing *in vitro*.^39^ The compound C1 was shown to reduce mature miR-377 levels in endothelial cells and restores VEGFA expression.^39^ Despite C1’s demonstrated selectivity and cellular activity, an atomic-level understanding of its interaction with pre-miR-377 remains unclear.

In this work, an integrative structural biology approach combining advanced multidimensional NMR spectroscopy with enhanced-sampling molecular dynamics simulations and Bayesian/maximum-entropy ensemble reweighting approaches was employed ^31,40^ to elucidate the atomic-level mechanism governing C1 recognition by pre-miR-377. This approach reveals how ligand binding redistributes pre-miR-377 conformational populations and identifies key molecular principles that govern selectivity. These findings provide mechanistic insights into RNA-small molecule recognition that can inform rational design strategies, establishing a framework for understanding recognition mechanisms across diverse RNA targets and advancing structure-based approaches in RNA-targeted therapeutics.

## RESULTS

Effective targeting of RNAs in general but especially pre-miR-377 with small molecules requires comprehensive characterization of its structure and associated conformational dynamics. High-resolution experimental techniques were integrated with advanced computational models to map the conformational landscape of pre-miR-377, both in its apo state and in complex with small molecule C1. This investigation focused particularly on the Dicer cleavage site and apical loop, two critical regions that govern recognition by the Dicer endonuclease complex and determine processing efficiency.

### NMR structural characterization reveals dynamic base-pairing patterns and conformational exchange in pre-miR-377

The first objective was to determine the exact 2D structure of pre-miR-377 and identify regions exhibiting dynamic behavior. To map the base-pairing network, ¹H-¹⁵N Correlation Optimized Spectroscopy (HNN-COSY) NMR experiments were conducted using fully ¹⁵N and ¹³C labeled RNA samples, which detect long-lived signals corresponding to base pairs stabilized at equilibrium (seconds to hours). The RNA construct contained two mutations over *wild type* sequence (U1G and A27C) to ensure successful synthesis and stabilization of the base of the stem. Site-specific assignments were achieved using selectively labeled samples at H3N3 and H1N1 positions of uridine and guanosine respectively, H8C8 and H2C2 positions of adenine, H8C8 position of guanosine and H6C6 position of uridine and cytosine. Additional selectively labeled samples included ^19^F at position 2’ and 5 of uridine. Selectively labeled bases were G1, G2, A4, U5, U6, C7, U10, U11, A13, U15, U16, A17, U18, G19, G22, A24, U25 and C26 achieving unambiguous assignments (**Figures 1, S1-S6**, and **Table S1**).

**Figure 1.**
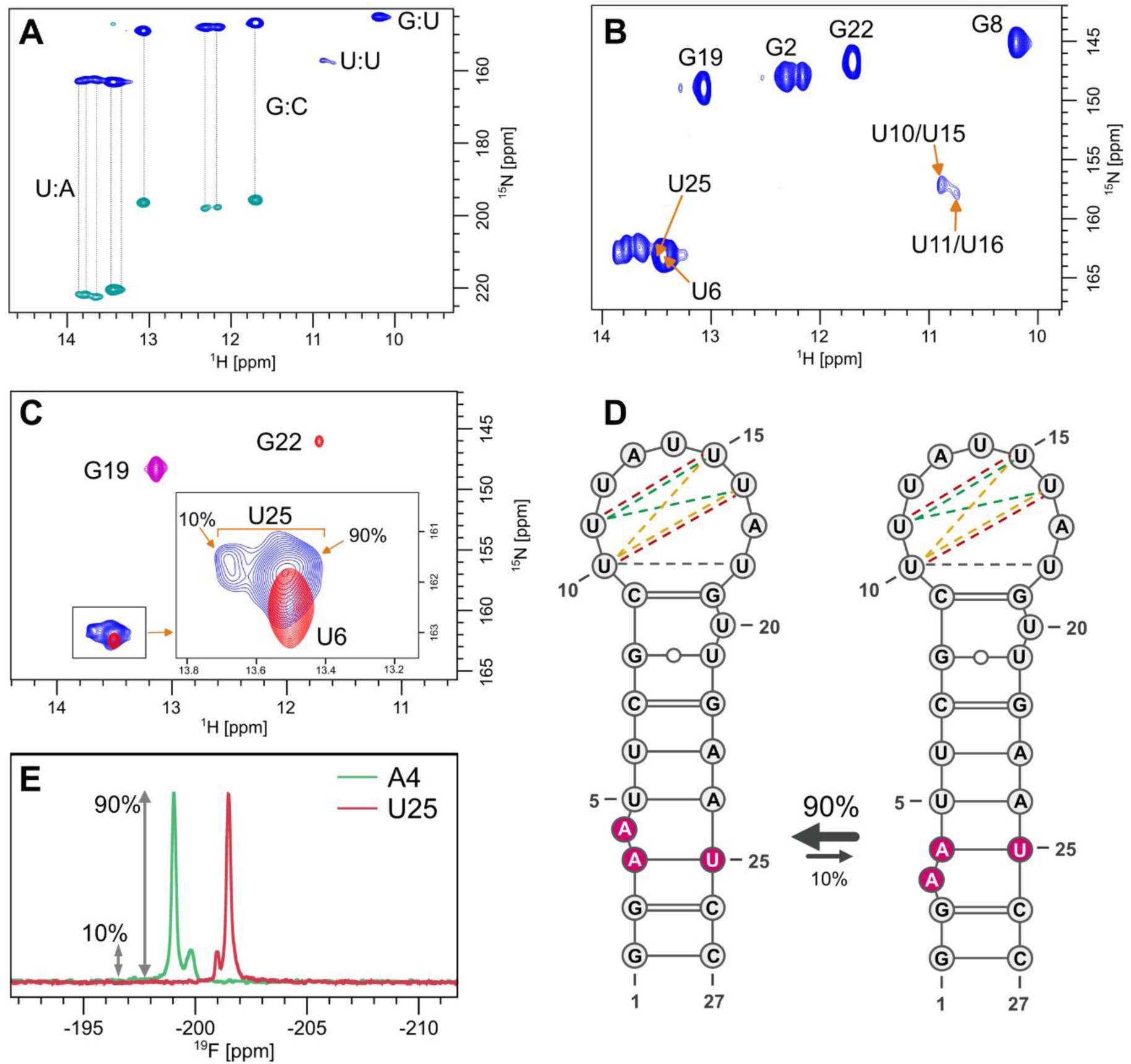
Secondary structure and conformational dynamics of pre-miR-377. **(A)** ¹H-¹⁵N HSQC spectrum of uniformly ¹⁵N-labeled pre-miR-377 showing imino proton correlations corresponding to Watson-Crick and non-canonical base pairs. Resonances are assigned based on sequential correlations and site-specific labeling. **(B)** ¹H-¹⁵N HSQC spectra of site-specifically labeled U10/U15 and U11/U16 pairs, confirming the presence of transient base pairing within the apical loop. **(C)** ¹H-¹⁵N HSQC spectrum of selectively labeled residues (U6, G19, G22, and U25) showing distinct resonances for U25 corresponding to two conformational states with ∼90:10 population ratio. **(D)** Secondary-structure representation of the two interconverting pre-miR-377 conformations, with arrows indicating the equilibrium between major (90%) and minor (10%) states. At the Dicer site, the shuffling of A3 and A4 with U25 results in the formation of two alternative states: A3-U25 and A4-U25 pairings. At the apical loop, transient and mutually exclusive base pairs are shown as dotted lines; each color indicates a set of base pairs that can form simultaneously. **(E)** ¹⁹F NMR spectra of 2′-fluoro-substituted A4 (green) and U25 (red) confirm the presence of two interconverting conformations with approximately 9:1 population ratio.

The NMR data revealed a well-defined stem structure with canonical Watson-Crick base pairs G1-C27, G2-C26, U5-A24, U6-A23, C7-G22, G8-U21, and C9-G19, while U20 forms a single-nucleotide bulge below the apical loop (**Figure 1A-B** and **D**). The Dicer cleavage site, however, exhibited remarkable structural heterogeneity. The ¹H-¹⁵N-SOFAST HMQC and HNN-COSY experiments on uniformly labeled and unlabeled samples revealed five distinct peaks in the U-A base pair region (13.5-15 ppm ¹H, 160-164 ppm ¹⁵N, **Figure 1B**). Two peaks were assigned to U6 (confirming the U6-A23 pair, **Figure 1C**) and U5 (conforming the U5-A24 pair, **Figure S5**), respectively, while two signals appearing in a 9:1 ratio were attributed to U25 via selective labeling (**Figure 1C**), suggesting that this nucleotide exhibits dynamic behavior by alternatively interacting with A3 and A4 at the Dicer site (**Figure 1D**). To characterize this conformational exchange, ¹⁹F labels were introduced at the 2′-ribose position of U25 and A4 and ^19^F NMR was performed to monitor backbone dynamics. These experiments confirmed two conformational states for both residues with a 9:1 population ratio (**Figures 1E**). Orthogonal ¹⁹F labeling at the 5-position of U25 further validated this behavior, revealing two populations (∼90% and ∼10%) with the major conformation displaying a sharp ¹⁹F signal characteristic of base pairing due to restricted mobility (**Figure S7**).^41^ These data suggest that U25 samples both paired and unpaired states with complementary population distributions: a 1:9 paired:unpaired ratio with A3 versus a 9:1 ratio with A4, indicating a dynamic adenosine bulge (A-bulge) at the Dicer site. Additional ¹⁹F analysis of U5 revealed further complexity, with this nucleotide predominantly paired (50%) while also sampling two additional conformations (30% and 20%, **Figure S7**), underscoring the complex conformational dynamics occurring the Dicer cleavage site of pre-miR-377.

Nonetheless, this analysis did account for the on remaining set of unassigned resonances in the U-A pairing region (13.5-15 ppm (¹H) and 160-164 ppm (¹⁵N); **Figure 1B**), suggesting these resonances may be due to uracil in the apical loop engaging in transient base pair interactions. To test this possibility, samples with ¹⁵N labels at H3N3 of U10, U11, U15, and U16 (**Figures S1** and **Table S1**) were designed. ¹H-¹⁵N-SOFAST HMQC experiments revealed complex H3N3 signals in the U-U pairing region (10-11 ppm),^42^ consistent with multiple non-canonical U-U interactions (**Figures 1B** and **S2**). Two resonances were observed for samples labeling U11 and U16, whereas a single major signal for U10 and U15 (**Figure S2C**). Because only the H3N3 signal can be observed in per U-U pair, the appearance of a single resonance indicated pairing with a unique partner, whereas two resonances indicate exchange between conformers. Together this pattern supports the presence of three mutually exclusive, interconverting apical-loop conformations (**Figure 1D**).

To further characterize the apical loop dynamics, we analyzed two additional ¹⁹F labeled constructs at U10 or U18. In both cases, ^19^F NMR spectra revealed two populations with approximate 60:40 ratios (**Figure S7**), which the major population exhibiting a sharp ¹⁹F signal characteristic of base-paired nucleotides.^41^ These results are consistent with ¹H-¹⁵N measurements and indicate that the apical lop region samples transient U-U base pair confirmations.

Our multidimensional NMR analyses define the secondary structure of pre-miR-377 and reveal a dynamic architecture at functionally important regions. The stem region is structurally stable and supported by well-defined base pairing, whereas the Dicer site displays pronounced conformational heterogeneity arising from a shuffling adenosine bulge that alternates between A3-U25 and A4-U25 pairings with a ∼9:1 population ratio. The apical loop exhibits additional complexity, with evidence for transient, non-canonical base-pair interactions involving U10, U11, U15, U16, and U18; however, the precise connectivity of these interactions remains difficult to resolve due to their short lifetimes and the intrinsic flexibility of this region.

### NMR relaxation analysis revealed fast nucleotide translational and rotational motions throughout the pre-miR-377

Having established the base-pairing network of pre-miR-377 at equilibrium on seconds-to-hours timescales, it was next investigated the faster molecular motions occurring on picosecond-to-millisecond timescales. These rapid dynamics encompass local translational and rotational fluctuations like inclination and tipping of bases, which are crucial for understanding site-specific RNA conformational plasticity. Notably, NMR relaxation measurements provide an ideal experimental approach for characterizing such molecular motions across multiple timescales with site-specific resolution.^43^ Thus, complementary R₁ (longitudinal) and R₂ (transverse, R₁ρ) relaxation experiments were employed on selectively labeled samples to map the dynamic landscape of pre-miR-377. These measurements probe distinct aspects of molecular motion: R₁ relaxation is primarily sensitive to fast local fluctuations on picosecond-nanosecond timescales (low R₁ corresponding to high relaxation time and fast motions), while R₂ relaxation reports on both rapid motions and slower conformational exchange processes occurring on microsecond-millisecond timescales (high R₂ indicating slow motions and/or conformational exchange). It is important to note that a spinlock of 2 kHz was used for the R₁ρ NMR experiment, meaning that only exchange contributions faster than 500 s⁻¹ could be detected. The systematic analysis revealed distinct mobility signatures throughout the RNA structure (**Figure 2**). Several nucleotides, including G8, A13, A17, G19, G22, A23, and A24, exhibited reduced R₁ values (1.8 ± 0.3 s⁻¹, **Figure 1A**) compared to the average, indicating enhanced picosecond-timescale flexibility. Conversely, loop region nucleotides U12, A13, U15, U16, and U17 showed lower R₂ values (80 ± 15 s⁻¹, **Figure 2B**), suggesting involvement in microsecond-millisecond motional processes.

**Figure 2.**
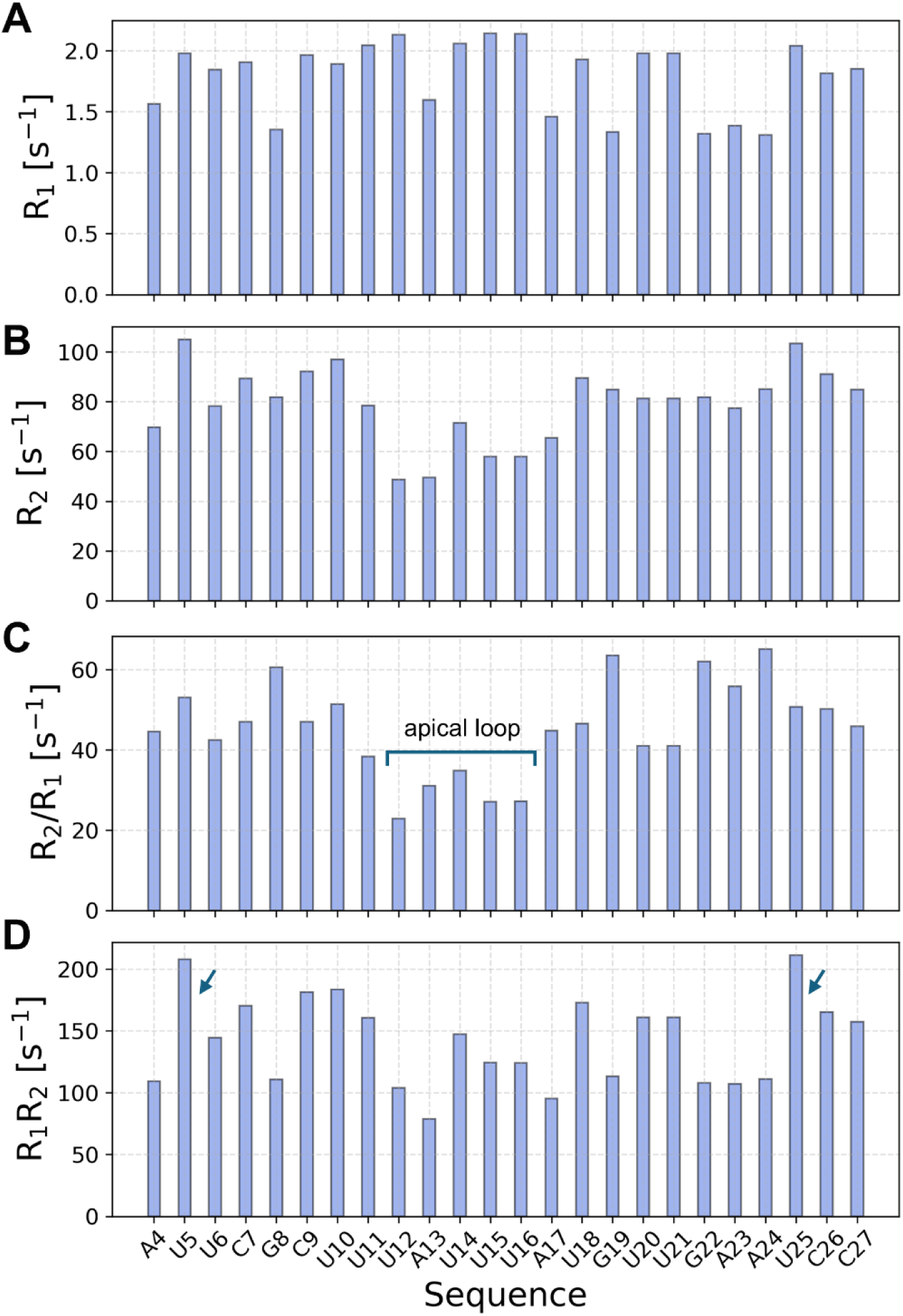
NMR relaxation analysis of pre-miR-377 as a function of nucleotide position. **(A)** *R*_1_ rates (s⁻¹). **(B)** *R*_2_ (*R*_1ρ_) rates (s⁻¹) measured under a 2 kHz spin-lock. **(C)** *R*_2_/*R*_1_ ratio across the sequence; nucleotides at the apical loop display lower ratios consistent with enhanced fast motions on the ps-ns timescale. **(D)** *R*_1_*.R*_2_ product; nucleotides with elevated values indicate μs-ms chemical exchange.

Moreover, the computed R₂/R₁ ratio provides insight into both local flexibility and chemical exchange phenomena.^44,45^ This analysis showed that apical loop nucleotides (U12, A13, U14, U15, U16) displayed significantly reduced R₂/R₁ ratios (45 ± 12, **Figure 2C**) relative to the stem regions, confirming their enhanced mobility. This analysis alone, however, cannot differentiate between intrinsic flexibility and conformational exchange and can be affected byanisotropic motions. For this reason, the R₁R₂ product was also computed that cancels out motional anisotropy^45^ and offers a more discriminating probe of the structural dynamics, since it is reduced by fast local motions but increased by slower conformational exchange processes.^45^ This R₁R₂ analysis revealed that U5 and U25 exhibit markedly elevated R₁R₂ values (140 ± 40, **Figure 2D**) compared to other nucleotides, providing strong evidence for microsecond-millisecond conformational exchange. This finding directly supports the earlier assignment of these residues as the source of the additional U-A cross-peaks observed in the 13.5-15 ppm (¹H) and 160-164 ppm (¹⁵N) spectral region, in line with the observation of two distinct U25 populations in ¹⁵N HMQC and ¹⁹F experiments **(Figures 1D, S7)**.

The comprehensive NMR relaxation analysis reveals a multi-timescale dynamic landscape within pre-miR-377. The apical loop exhibits rapid ps-ns motions, while U5 and U25 at the Dicer site undergo µs-ms conformational exchange compatible with alternative base-pairing states.

### Atomistic 3D structural characterization of pre-miR-377 through NMR-informed *in silico* modeling

Following identification of the secondary structure and flexible regions, we characterized the three-dimensional conformational ensemble of pre-miR-377 at atomistic resolution using an integrative modeling approach. We first used the FARFAR2 fragment assembly method to generate 750 three-dimensional models consistent with the most populated NMR-defined secondary structure (**Figure 1D**), enforcing only stable base-pair interactions and excluding the transient and interconverting interactions observed in the apical loop. The top-ranked FARFAR2 model was then used to initiate extensive enhanced-sampling molecular dynamics (MD) simulations to explore the conformational landscape. Specifically, we performed Hamiltonian replica exchange simulations combined with well-tempered metadynamics, using the RNA radius of gyration as the collective variable, and applied two state-of-the-art RNA force fields (χOL3 with backbone phosphate modification^46–49^ and DESRES^50^) for a total simulation time of 6 μs (**Figure S8A-C**).

To improve *in silico* accuracy and agreement with experimental data, we applied a Bayesian/maximum-entropy reweighting strategy previously validated for flexible RNAs.^31,51,52^ We collected 2D and 3D NOESY spectra in H₂O and D₂O, yielding an average of seven restraints per nucleotide (175 non-redundant restraints; **Table S2**, **Figure S9**). These restraints report on proton-proton distances involving imino, amino, aromatic, and ribose protons.^43^ For each modeled conformation, inter-proton distances were back-calculated and weighted according to agreement with the experimental data (**Figures S10-S11**).

Reweighting substantially improved agreement between computational ensembles and NMR restraints across all methods, as reflected by reduced χ² values (**Figure 3A**). Molecular dynamics simulations produced ensembles with strong agreement independent of force field (χ² = 1.9 for χOL3 and χ² = 1.3 for DESRES). In contrast, FARFAR2 showed the weakest agreement (χ² = 3.5; **Figure 3A**), consistent with prior reports that fragment-assembly approaches may insufficiently sample RNA conformational space.^53^

**Figure 3.**
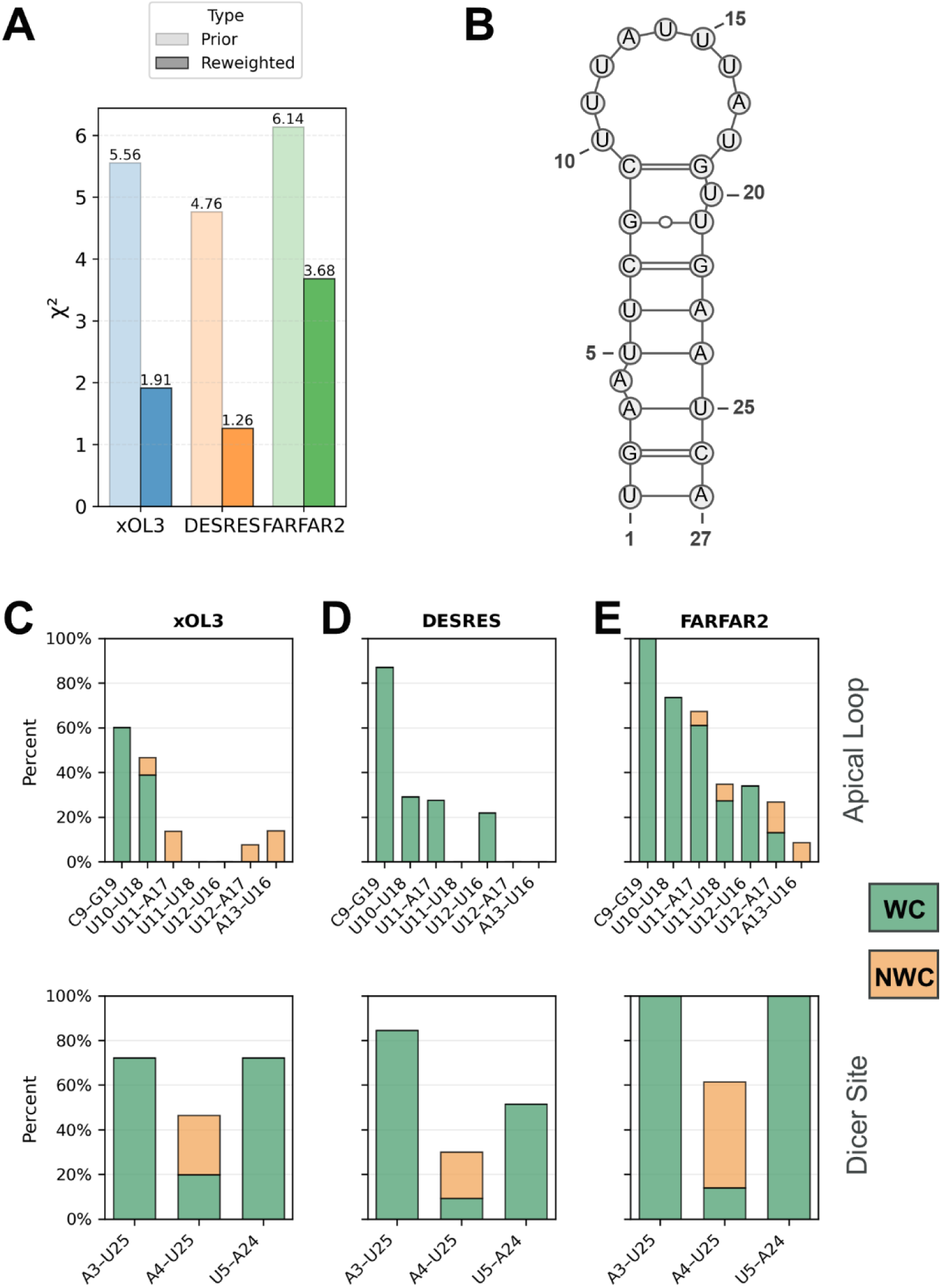
Evaluation of *in silico* pre-miR-377 structural models against NMR restraints. **(A)** Comparison of χ² values between RNA *in silico* structural models and experimental NMR proton-proton distance restraints for three modeling approaches: Molecular Dynamics simulations using χOL3 (blue) and DESRES (orange) force fields, and de novo fragment assembly algorithm implemented in FARFAR2 (green). Bars represent χ² values for prior (light) and reweighted (dark) ensembles. Lower χ^2^ values show better agreement with experimental data. **(B)** Reference 2D structure of the major pre-miR-377 conformation detected by NMR. **(C-E)** Base-pair composition and distribution in the apical loop (top) and Dicer-site (bottom) regions derived from the (C) χOL3, (D) DESRES, and (E) FARFAR2 ensembles. Base pairs are classified as Watson-Crick (WC, green) or non-Watson-Crick (NWC, orange), as shown in the legend. The bar heights represent the percentage population of each base-pair type within the respective regions.

The largest differences between ensembles were observed in regions identified by NMR as flexible, namely the apical loop and the Dicer site. In the apical loop, both MD ensembles adopted largely unstructured conformations with few persistent base-pair interactions (**Figures 3C-D**). In the χOL3 ensemble, base pairing was primarily limited to nucleotides near the stem, including C9-G19 (58% population) and U10-U18 (38%), with additional transient non-canonical interactions involving U11, U12, and U16 (**Figure 3C**). These features partially agree with the experimental data (**Figure 1**).

In contrast, FARFAR2 predicted multiple stable loop interactions (**Figure 3E**), including C9-G19 (85%), U10-U18 (74%), U11-A17 (67%), U11-U18 (35%), U12-U16 (34%), and U12-A17 (27%), which is inconsistent with the high flexibility observed by NMR in this region. Notably, none of the computational ensembles adequately sampled the non-canonical U10/U15 and U11/U16 interactions detected by NMR (**Figure S2C**), highlighting persistent challenges in modeling highly dynamic RNA regions, consistent with the current state of the field.^54^

At the Dicer site, both MD ensembles reproduced the NMR-observed shuffling A-bulge (**Figures 3C-D**). In the χOL3 ensemble, A3-U25 and U5-A24 formed Watson-Crick base pairs in 72% of the population, whereas A4-U25 Watson-Crick pairing occurred in only 20%, with an additional 26% adopting non-canonical interactions (**Figure 3C**). These results agree with NMR data showing dynamic exchange among A3/A4-U25 and U5-A24 base pairs as key contributors to conformational heterogeneity at the Dicer A-bulge. By contrast, FARFAR2 failed to capture this flexibility, maintaining persistent Watson-Crick A3-U25 and U5-A24 base pairs in 100% of the ensemble, with A4 interacting with U25 only when U25 was disengaged from A3 (**Figure 3E**).

Collectively, these results demonstrate that NMR-reweighted MD simulations show substantially better agreement with experimental data than FARFAR2 and more accurately capture the structural dynamics of both the apical loop and the Dicer site.

### A-bulge shuffling induces the spontaneous formation of two RNA triples at the Dicer site of pre-miR-377

Because FARFAR2 showed the weakest agreement with the NMR data, subsequent analyses focused exclusively on MD-derived ensembles. We performed structural clustering to identify the most frequently sampled three-dimensional conformations within the pre-miR-377 MD ensembles (**Figures 4 and S12–S13**).

**Figure 4.**
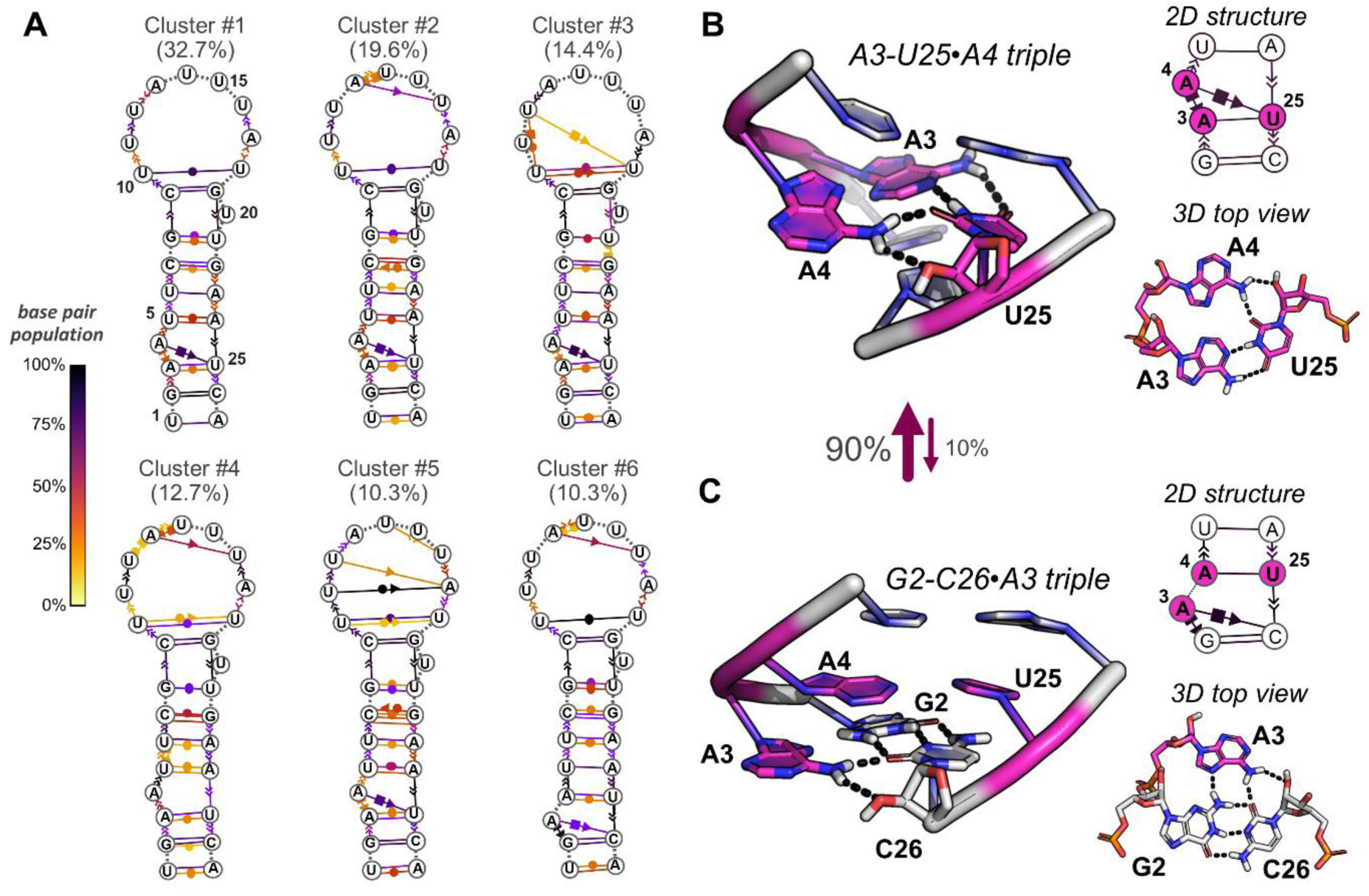
Structural clustering and base-triple formation in pre-miR-377. **(A)** Secondary structure of the six major clusters identified within the χOL3 ensemble. The percentage shown for each cluster indicates its relative population among all clustered structures. Base-pair interactions are annotated using the Leontis-Westhof classification and colored according to their population density within the cluster (yellow to black), with black indicating higher population. **(B)** Representative 2D and 3D structure of the A3-U25•A4 triple, the dominant conformation observed at the Dicer site of pre-miR-377 (90% population). **(C)** Representative 2D and 3D structure of the alternative G2-C26•A3 triple conformation observed in approximately 10% of clustered structures. In both panel B and C, the nucleotides involved in the A-bulge shuffling, i.e., A3, A4, and U25, are shown in magenta, while hydrogen bonds are shown as black dotted lines.

Clustering of the χOL3 ensemble identified six distinct conformational clusters (**Figure 4A**), all of which maintained a largely base-paired stem region, consistent with NMR observations. In contrast, the apical loop exhibited substantial heterogeneity across clusters, with differences in both base-pairing and stacking interactions. The most populated cluster (cluster #1; 32.7%) featured C9-G19 and U10-U18 base pairs at the base of the loop, accompanied by continuous stacking from C9 to A13 and from U16 to U18. A *cis* sugar-sugar A13-U16 interaction was observed in clusters #2, #4, and #6, each sampling distinct loop conformations. For example, cluster #2 additionally displayed a *cis* sugar-Hoogsteen interaction between A13 and U14, whereas cluster #4 sampled a *cis* interaction between the Watson-Crick edge of U12 and the Hoogsteen edge of A13, underscoring the dynamic and structurally diverse nature of the apical loop.

Notably, clustering of the χOL3 ensemble revealed two mutually exclusive A-bulge conformations at the Dicer site, defined by alternative A3-U25 and A4-U25 base pairing. The A3-U25 conformation dominated clusters #1-#5 (∼90% of the ensemble), while the A4-U25 conformation was confined to cluster #6 (∼10%; **Figure 4A**). This ∼9:1 population distribution was consistent with the A-bulge shuffling observed by both ^1^H-^15^N SOFAST-HMQC and ^19^F NMR experiments (**Figures 1C-E**). Beyond reproducing the experimental populations, structural analysis revealed that this base-pair exchange gives rise to two distinct RNA triple structural motifs. When A3 pairs with U25, A4 primarily forms a sugar-Hoogsteen interaction with U25, aligning in the same plane as the A3-U25 pair to generate an A3-U25•A4 triple in clusters #1, #2, #3, and #5 (**Figure 4B**). This configuration is consistent with additional U-A resonances observed between 13.5-15 ppm (^1^H) and 160-164 ppm (^15^N) in the NMR spectra (**Figure 1B**). In a less populated state (cluster #4; 12.7%), A3-U25 pairing persists while A4 is fully bulged out (**Figure 4A**). Conversely, in cluster #6, A4 pairs with U25, and A3 engages the downstream G2-C26 pair via a Hoogsteen interaction, forming a G2-C26•A3 triple (**Figure 4C**).

In contrast, clustering of the ensemble generated with the DESRES force field returned two clusters both characterized by one single conformation of the Dicer site involving the A3-U25, differing solely in A4 positioning between the two clusters: forming an A3-U25•A4 triple in cluster #1 (**Figure S13A**), and stacking with A3 in cluster #2 (**Figure S13B**). Notably, this single Dicer site conformation characterized by the A3-U25 base pair is in contrast with NMR data showing the formation of both the A3-U25 and A4-U25 pairs in a 9:1 ratio. Moreover, the NMR-detected G8-U21 pair was sparsely sampled in both clusters, which were characterized by numerous and steady interactions at the apical loop involving U10-U18, U11-A17, U12-U16, and A13-U15 contacts (**Figure S13**) – inconsistent with NMR data showing only transient interactions at the apical loop.

Collectively, these data demonstrate that the ensemble generated with the χOL3 force field fits accurately most of the NMR data, revealing that the Dicer site can sample two major conformational states due to U25 shuffling between A3 and A4 that can alternatively pair with U25 to form distinct RNA triples.

### Multidimensional NMR spectroscopy confirmed specific binding of small molecule C1 to the Dicer cleavage site of pre-miR-377

The integrated structural analysis of pre-miR-377 apo dynamics prompted us to investigate whether and how small molecule binding affects the structural ensemble of this system. A compound was identified, dubbed C1, that binds to the Dicer processing site of pre-miR-377, modulating its maturation and promoting increased VEGFA expression.^39^ Building upon these initial results showing C1 binding to pre-miR-377, it was further studied how C1 interacts with its target with mono- and two-dimensional NMR spectroscopy (**Figure 5**, and **S14**).

**Figure 5.**
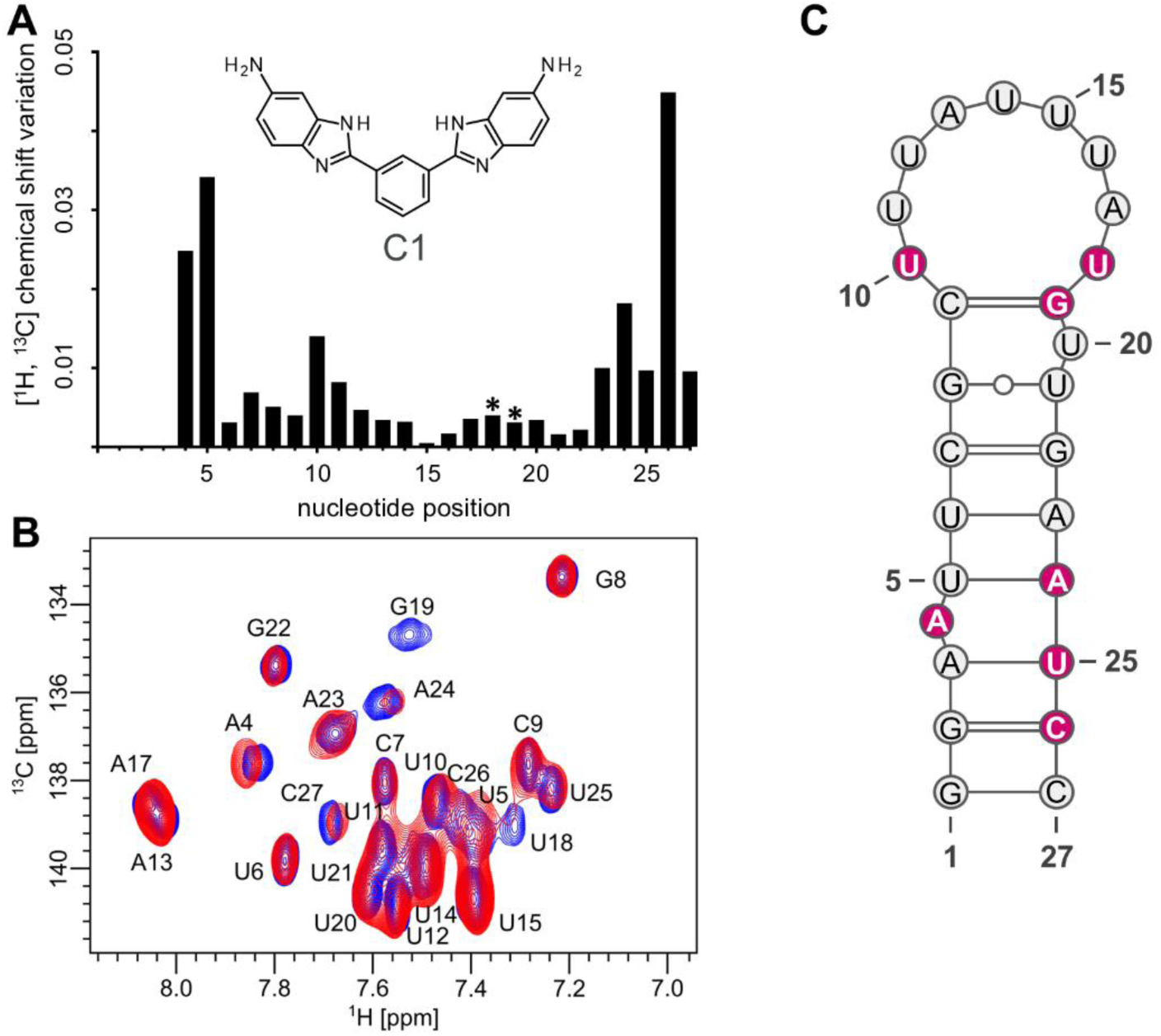
NMR characterization of C1 binding to pre-miR-377. **(A)** ^1^H-^13^C chemical shift perturbations (CSPs) for assigned aromatic resonances comparing two concentrations of small molecule C1, i.e., 0 vs 100 µM; largest CSPs map to A4, U5, U10, A24, and C26 at the Dicer site. The chemical structure of the small molecule C1 is also shown. The asterisk key (*) indicates that these residues broaden beyond detection due to intermediate exchange in the NMR timescale. Hence, CSP for these nucleotides are not quantified. **(B)** Overlay of ^1^H-^13^C HSQC/HMQC spectra in the H8C8 and H6C6 regions for pre-miR-377 at 0 µM (blue) and 100 µM C1; expansions highlight A4 (fast-exchange shifts) and A24 (line broadening indicative of slow-to-intermediate exchange). **(C)** Secondary-structure of the major conformation of pre-miR-377 detected by NMR with nucleotides exhibiting the largest chemical shift from both panel A and B highlighted in magenta, delineating the C1 binding site at the Dicer cleavage region.

First, 1D NMR Carr-Purcell-Meiboom-Gill (CPMG T2) experiments^55^ were used to monitor spectral changes in C1 upon the addition of pre-miR-377. These experiments revealed substantial decreases in peak intensity of compound C1 upon increasing concentrations of pre-miR-377 (**Figure S14**), indicating a direct interaction between the two molecules.

To pinpoint the binding site and characterize the nature of the interaction, we carried out 2D NMR Heteronuclear Multiple Quantum Correlation ^1^H-^13^C experiments^43^ on site-specifically labeled **(Figure S1**) pre-miR-377 with a saturating concentration of C1. Chemical shift perturbations were observed for nucleotides A4, U10, U18, G19, A24, U25, and C26 (**Figure 5A-C**). Notably, this region mainly comprises the Dicer recognition site of pre-miR-377, providing a molecular rationale for the interference of C1 with Dicer-mediated cleavage.^39^

Furthermore, the bases experiencing chemical shift perturbations generally exhibited decreased signal intensity upon increasing concentrations of C1. This behavior is consistent with binding in an intermediate exchange time regime, corresponding to low micromolar affinity, which corroborates previously measured affinities for this interaction (K_d_ = 29 ± 0.3 µM).^39,43^ Due to the broadening of NMR peak intensity on increasing concentration of C1, however, we were unable to determine the affinities directly from these changes.

Thus, binding studies reveal that C1 specifically associates with the Dicer site of pre-miR-377, resulting in observable conformational changes in the RNA. This interaction likely disrupts or alters Dicer site conformation, providing a mechanistic basis for the observed modulation of miRNA maturation and subsequent increase in VEGFA expression.^39^

### Small molecule C1 selectively binds pre-miR-377 through a pathway-dependent mechanism

To elucidate the C1-pre-miR-377 binding mechanism at atomic detail, we conducted comprehensive out-of-equilibrium molecular dynamics (MD) simulations to enhance sampling of binding events at the Dicer site. We used two NMR-validated pre-miR-377 structures representing the major (90%) and minor (10%) Dicer site populations from χOL3 sampling without C1, dubbed A3-U25 and A4-U25 based on their mutually exclusive base pairing patterns. Each conformation was simulated with C1 initially positioned at 12 Å from the Dicer site at either minor groove (A3-U25_min, A4-U25_min) or major groove (A3-U25_mj, A4-U25_mj) to explore alternative binding pathways. Based on experimental evidence that C1 cannot bind a pre-miR-377 mutant,^39^ we included this variant as negative control. The mutant contains one uracil insertion between A24 and U25 that pairs with A4, eliminating A-bulge shuffling and locking the Dicer site in a single helical conformation. Also, the mutant was simulated with C1 approaching both groove orientations (mutant_min and mutant_mj). For each system, we performed 200 steered MD simulations biasing the C1-Dicer distance with carefully designed protocols ensuring equilibrium sampling (see Methods), for a total of 6 µs simulation time.

We monitored the distance between C1 and the Dicer site in all simulations and observed that the wild-type pre-miR-377 recognizes C1 regardless of initial conformation, but it exhibits a strong preference for minor groove approach orientations, achieving substantially closer binding geometries (**Figure 6A**). Indeed, the A3-U25 systems yielded median C1-Dicer site distances of 3.4 Å versus 5.7 Å for minor and major groove approaches, respectively, similarly to what observed for the A4-U25 systems, which showed corresponding distances of 4.3 Å and 6.4 Å. Notably, both mutant systems consistently maintained distances ≥ 5.7 Å regardless of approach orientation, confirming C1’s inability to bind the rigid helical Dicer site architecture of the mutant.^39^

**Figure 6.**
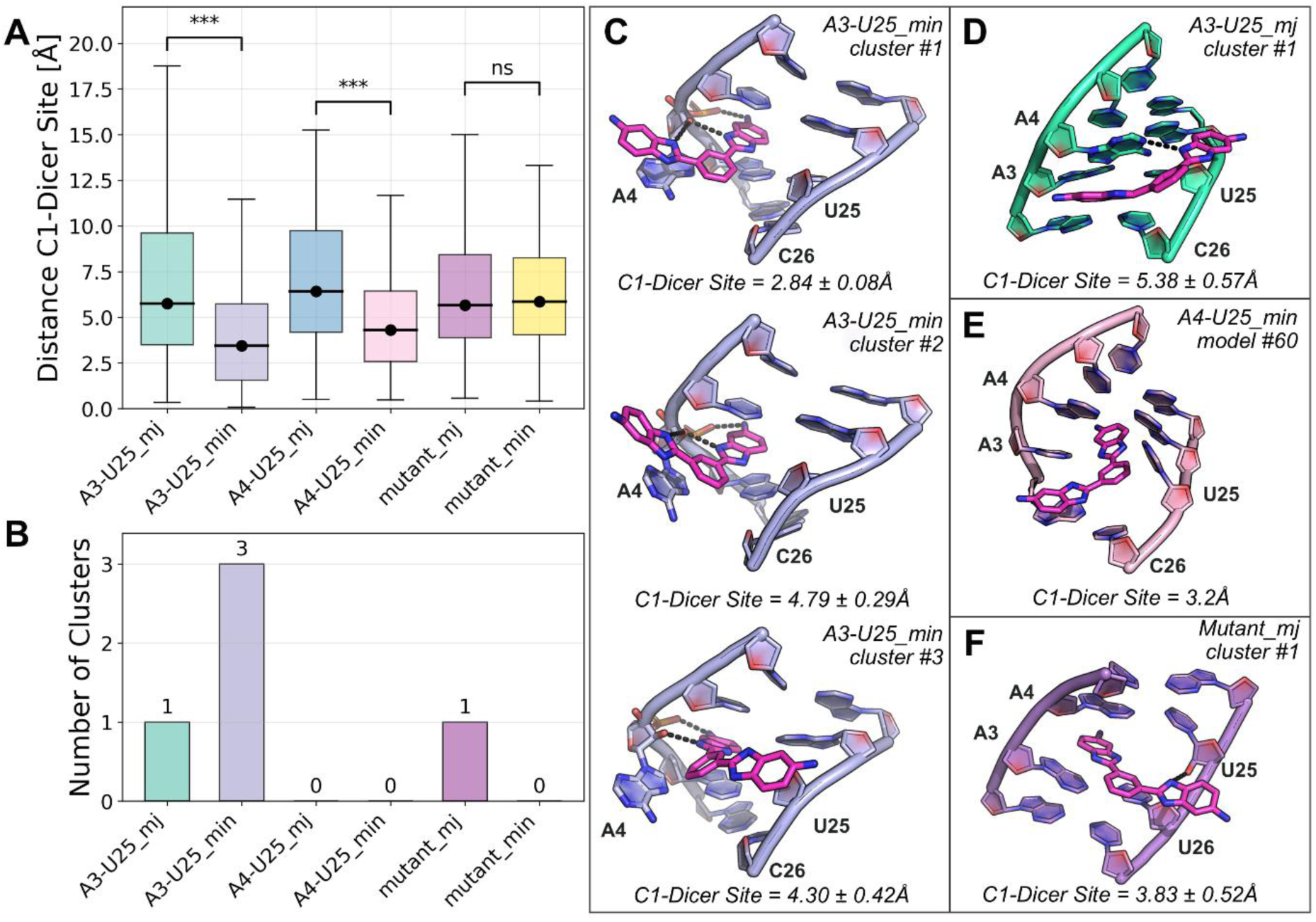
Sampling and conformational analysis of C1-Dicer site interactions across pre-miR-377 variants. **(A)** Boxplots showing the distribution of C1-Dicer site distances for wild-type (A11-U33, A12-U33) and mutant pre-miR-377 variants sampled from major (mj) and minor (min) groove pathways. Black dots represent median values, and error bars indicate the range of sampled distances. Statistical significance is denoted as triple stars when p < 0.001; ns, not significant. **(B)** Number of conformational clusters identified for each system, defined by structural similarity of C1 and surrounding Dicer-site nucleotides (RMSD < 2.5 Å within a cluster). **(C–F)** Representative structures of the dominant conformational clusters for each pre-miR-377 system. Nucleotides at or near the Dicer site are shown as cartoons (colored according to panel A), and the small molecule C1 is depicted as magenta sticks. Hydrogen-bonding interactions are indicated as black dotted lines, where relevant. Average C1-Dicer site distances with their standard deviations are annotated for each cluster or model.

Cluster analysis of the accumulated trajectories provided detailed insights into the binding mode diversity across systems (**Figure 6B**). The A3-U25_min system generated three distinct yet structurally related clusters, indicating a conserved binding core with limited conformational variability. Across all three clusters, C1 binding destabilized the A3-U25•A4 triple conformation at the Dicer site, with A4 consistently displaced from the A3-U25 base pair plane. The first two clusters exhibited similar binding modes, differing only in variations of the stacking plane orientation (**Figure 6C**). In these configurations, one of the two symmetric benzimidazole moiety of C1 is sandwiched between U5 and A3, while the other half of C1 engages in an additional stacking with A4, and establishes three hydrogen bonds with A4’s ribose and phosphate backbone. The third cluster maintained this core interaction network but featured a 180° rotation of the phenylbenzimidazole moiety around its plane, eliminating the A4 stacking interaction while partially preserving the hydrogen bonding pattern. In contrast, A3-U25_mj yielded only a single cluster characterized by markedly different binding characteristics (**Figure 6D**). Here, the Dicer site was stabilized in a A3-U25•A4 triple conformation, and C1 remained completely solvent-exposed, showing no stacking interactions and only a single hydrogen bond with A4 at the Dicer site, confirming the structural liability of binding modes sampled from major groove paths (**Figure 6A**).

Compound C1 binding to the A4-U25 conformation revealed fundamentally different recognition behavior, with no identifiable clusters regardless of pathway (**Figure 6B**). While C1 rarely engaged from the major groove (median distance 6.4 Å), minor groove binding occurred but profoundly destabilized the base-pairing network at the Dicer site **(Figure 6E**). This extensive perturbation generated multiple binding modes too heterogeneous to cluster, suggesting this less populated A4-U25 conformation (10% occupancy) cannot accommodate C1 through a unique binding pose.

The mutant showed opposite clustering patterns: only major groove approach (mutant_mj) yielded clusterable binding modes, suggesting fundamentally different recognition mechanisms between helical and bulge-containing structures **(Figure 6F**). However, this cluster showed C1 only partially intercalated, establishing one hydrogen bond with U25’s sugar while remaining largely solvent-exposed, resembling the A3-U25_mj binding mode. The convergence on similar binding modes despite different RNA architectures suggests major groove binding generally favors highly solvent exposed poses, weak interactions lacking structural requirements for stable recognition.

Collectively, these simulations demonstrate that C1 selectively recognizes wild-type pre-miR-377 through pathway-dependent mechanisms strongly favoring minor groove approaches. Binding involves substantial Dicer site reorganization relative to apo pre-miR-377, consistent with NMR-observed chemical shift variations.

### C1 binding shifts pre-miR-377 conformational equilibrium towards Dicer-incompatible states

To characterize the binding energetics underlying C1 recognition, we performed Hamiltonian replica exchange MD (H-REMD) simulations coupled with well-tempered metadynamics. Starting from C1-unbound models, these simulations explored C1 binding from the minor groove to three RNA systems: wild-type Dicer site conformations A3-U25 and A4-U25, and the mutant variant. The distance between C1 and the Dicer site served as the collective variable, while allowing the Dicer site to freely explore its conformational landscape and capture the full spectrum of binding-induced structural changes.

The H-REMD simulations showed distinct thermodynamic signatures for each system (**Figure 7A**). Both wild type systems exhibited well-defined free energy minima corresponding to C1-bound states at distances between 1-4 Å from the Dicer site. In contrast, the mutant showed a monotonically increasing free energy profile with decreasing C1-Dicer site distance, indicating that C1-bound conformations are thermodynamically disfavored by approximately 4 kcal·mol^-1^ relative to unbound states (**Figure 7A**). This quantitative analysis provides direct thermodynamic evidence for the complete loss of binding affinity upon mutation of pre-miR-377 as experimentally demonstrated by Haniff et al.^39^

**Figure 7.**
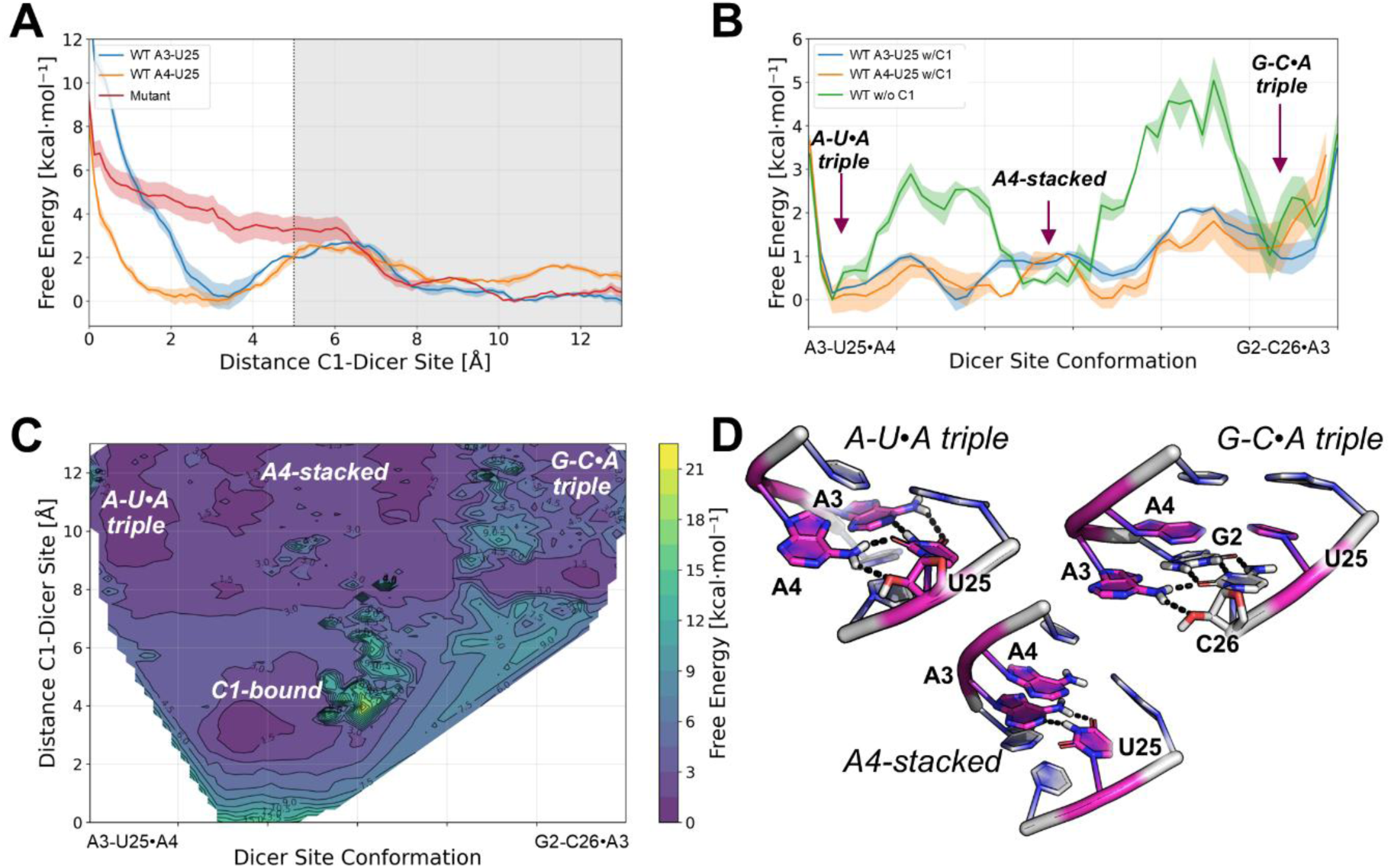
Small molecule C1 shifts the conformational equilibrium of pre-miR377. **(A)** Free energy profiles as a function of the C1-Dicer site distance for wild-type (A11-U33, A12-U33 input conformations) and mutant pre-miR-377 variants. Errors are reported as shaded areas. The vertical dashed line marks the boundary between bound and unbound states. **(B)** Free energy profiles along a path collective variable describing the Dicer site conformational transitions both in the presence (blue and orange lines) and absence of C1 (green line). Characteristic conformations corresponding to A-U•A, A4-stacked, and C-G•A triple states are indicated. The presence of C1 reduces the barriers between different conformations of the Dicer site. **(C)** Two-dimensional free energy surface showing the coupling between the C1-Dicer site distance and Dicer site conformation for the wild-type system. Distinct minima corresponding to the C1-bound, A-U•A triple, A4-stacked, and C-G•A triple conformations are labeled. **(D)** Representative structures of key conformational states. RNA is shown as grey cartoon, while nucleotides responsible for the A-bulge shuffling (A3, A4, and U25) are highlighted as magenta sticks. Hydrogen bonds are shown as black dotted lines. Representatives C1-bound states are shown in Figure 6C.

The comparative analysis of *wild type* systems revealed additional mechanistic insights. While both A3-U25 and A4-U25 conformations exhibited similar estimated binding barriers of approximately 2 kcal·mol^-1^, the free energy minimum corresponding to the C1-bound state was notably broader and shallower for the A4-U25 system. This observation suggests that C1 explores multiple structurally diverse but energetically similar binding modes when interacting with A4-U25 conformations, consistent with the heterogeneous binding behavior and absence of clusterable conformations observed in our orthogonal steered MD simulations for the same system (**Figure 6B**).

To understand how C1 binding influences the structural dynamics of its RNA target, we monitored Dicer-site conformational transitions during H-REMD using a path collective variable defined over nucleotides G2-U5 and A23-C26. The path was built between the two experimentally supported reference states; the A3-U25•A4 triple (cluster #1; **Figure 6C**) and the G2-C26•A3 triple (cluster #6; **Figure 6C**), and discretized into further 13 intermediate nodes uniformly spaced in eRMSD. This allowed us to quantify similarity to each state and track C1-induced conformational shifts along the conformational coordinate. In the apo state, the free energy landscape showed a clear preference for the A3-U25•A4 triple conformation, which represents the global minimum (**Figure 7B**). This stable state is separated from the alternative G2-C26•A3 triple by multiple barriers, with the highest reaching approximately 5 kcal·mol^-1^. An intermediate metastable state, characterized by an unpaired A4 extending toward the bulk solvent, appears as a local minimum separated from the A4•A3-U25 triple state by about 3 kcal·mol^-1^.

Most remarkably, C1 binding altered this conformational landscape. The presence of the small molecule in both A3-U25 and A4-U25 systems significantly flattened the free energy profile, reducing the barriers separating the two triple conformations by more than half (**Figure 7B**). This barrier reduction facilitated access to metastable states, resulting in a more uniform distribution of conformational states. Specifically, our simulations showed that C1 preferentially stabilizes intermediate conformations that correspond to partially disordered triple structures, rather than binding to the well-defined triple conformations themselves. This mechanism suggests that C1 recognition involves the formation of dynamic, partially structured states followed by stabilization of these otherwise transient conformations.

To provide a better visualization of this mechanism, we generated a two-dimensional free energy surface plot by combining the conformational coordinate on the X axes and the distance between C1 and the Dicer site on the Y axes (**Figure 7C**). The energy minimum corresponding to the C1 binding (C1-Dicer site distance is ∼3Å), occurs when the Dicer site adopts intermediate conformations between the two reference triples (**Figure 7C)**, confirming that optimal small molecule recognition requires departure from the stable triple structures.

Collectively, these data reveal that C1 achieves selectivity for wild-type pre-miR-377 through preferential recognition of conformationally dynamic bulge structures, while the rigid helical architecture of the mutant prevents productive binding interactions. Through a two-step mechanism, C1 first recognizes triple structures and then shifts the pre-miR-377 Dicer site conformational equilibrium.

## DISCUSSION

Targeting precursor microRNAs (pre-miRNAs) with small molecules represents a powerful but underdeveloped therapeutic strategy for diseases in which microRNA dysregulation drives pathology. A fundamental obstacle has been the intrinsic conformational plasticity of pre-miRNAs that complicates both structural characterization and rational ligand design. Unlike proteins, where stable folds often define druggable pockets, pre-miRNAs populate heterogeneous ensembles that interconvert on multiple timescales. As a result, ligand recognition cannot be understood from static structures alone but instead emerges from the interplay between RNA dynamics, binding pathways, and ligand-induced reorganization.

Here, we combined multidimensional NMR spectroscopy with extensive enhanced-sampling molecular dynamics (∼20 μs) and integrative ensemble refinement to define, at atomistic resolution, how the small-molecule inhibitor C1 selectively recognizes pre-miR-377, a therapeutically relevant target implicated in cardiovascular disease. This integrative framework reveals a two-step recognition mechanism in which C1 first engages a dynamically exchanging adenosine bulge (A-bulge) at the Dicer site through pathway-dependent minor groove access and subsequently reshapes the conformational ensemble toward states that are incompatible with productive Dicer processing. Importantly, this mechanism does not rely on recognition of a single static structure but instead exploits the RNA’s intrinsic ability to sample alternative architectures.

Site-resolved NMR measurements establish that pre-miR-377 is best described as a modular RNA with sharply distinct dynamic regimes (**Figure 1**). The stem region remains structurally robust, dominated by canonical Watson-Crick base pairs, whereas the Dicer site exhibits pronounced conformational exchange driven by shuffling of A3 and A4 in pairing with U25 at a ∼9:1 ratio (**Figure 1C-D**). This shuffling was independently confirmed using selective ^1^H-^15^N labeling and orthogonal ^19^F probes (**Figure 1E, S7**), while elevated R₁R₂ relaxation rate products for U5 and U25 revealed μs-ms exchange processes consistent with multiple interconverting conformations (**Figure 2D**). In contrast, the apical loop samples only transient and weakly populated non-canonical interactions accompanied by fast ps-ns motions, highlighting the technical limitations of resolving complete connectivity in highly dynamic RNA regions using NMR alone. Although additional approaches such as residual dipolar couplings, paramagnetic relaxation enhancement, or chemical exchange saturation transfer could further refine these regions, such experiments lie beyond the scope of the present work.

To translate these experimental observables into atomistic structural models, we generated ensembles using both fragment assembly (FARFAR2) and enhanced-sampling MD simulations, followed by Bayesian/maximum-entropy reweighting against NOE-derived distance restraints. MD-derived ensembles showed substantially improved agreement with NMR data relative to fragment assembly approaches (**Figure 3A**), emphasizing that conformational sampling, not model generation alone, is the dominant limitation for flexible RNAs. Simulations using the χOL3 force field quantitatively reproduced the experimentally observed A-bulge shuffling and revealed its structural consequences: the formation of two mutually exclusive RNA triple motifs at the Dicer site (**Figure 4A**).

When A3 pairs with U25 (∼90% population), A4 adopts a *cis* Hoogsteen-sugar interaction, forming an A3-U25•A4 triple (**Figure 4B**). Conversely, when A4 pairs with U25 (∼10%), A3 redirects to the downstream G2-C26 base pair via a Hoogsteen interaction, yielding a G2-C26•A3 triple (**Figure 4C**). The close quantitative agreement between experimental and computational population distributions (9:1) validates these assignments and demonstrates that ensemble-resolved modeling is essential for capturing functionally relevant RNA states that are invisible to static representations. Notably, while MD enabled detailed atomistic insight into the Dicer site, neither tested force field fully captured all NMR-detected interactions in the apical loop (**Figure 3A-D, S13**), underscoring the continuing need for experimental refinement when modeling highly dynamic RNA motifs.

Strikingly, the RNA triple motifs identified at the pre-miR-377 Dicer site are not idiosyncratic but instead correspond to evolutionarily conserved architectural elements found across diverse functional RNAs. Mining the RNA triple database^56^ revealed that both AUA- and CGA-type triples observed here recur in group I and II introns, ribosomal RNAs, riboswitches (ydaO), RNA aptamers (Chili), and CRISPR RNAs, including the Cas13b crRNA (**Table S3, Figure S15**). In the Cas13b system, a structurally analogous G128-C178•A129 triple directly contacts the RNase recognition site and contributes to RNA processing efficiency.^57^ This parallel raises the possibility that dynamic RNA triples at bulge regions represent a convergently evolved solution for regulating RNase engagement across biological systems, as has been observed for other RNA-processing enzymes such as Cas proteins and RNase H1.^58,59^ Whether the pre-miR-377 triples directly participate in Dicer recognition remains to be experimentally validated; however, structural superposition of both apo triple conformations onto the human Dicer-RNA complex (PDB ID: 7XW2) reveals steric clashes with the catalytically essential Arg1855 residue (**Figure S16B-D**), suggesting that either RNA or enzyme reorganization, or both, would be required for productive processing.

Against this structural backdrop, enhanced-sampling MD simulations of ligand binding reveal how C1 exploits RNA dynamics to achieve selective recognition (**Figure 6**). Productive binding occurred exclusively through minor groove trajectories, where C1 intercalates between U5 and A3 and engages in cooperative stacking and backbone hydrogen-bonding interactions (**Figure 6A and C**). Major groove approaches failed to form stable complexes (**Figure 6D-F**), reflecting fundamental features of RNA geometry: bulge-containing minor grooves provide confined, partially dehydrated environments that accommodate planar aromatic scaffolds while presenting accessible phosphate and sugar functionalities, whereas major grooves are sterically congested and highly solvated. This mirrors long-established principles in DNA recognition, where minor groove binders exhibit enhanced selectivity,^60,61^ and argues that groove accessibility should be treated as a first-order design parameter for RNA-targeted ligands.

Critically, C1 does not merely bind a pre-existing conformation but actively reshapes the RNA ensemble. Upon engagement, C1 disrupts native triple architectures by displacing A4 from the A3-U25 plane, thereby altering the geometry of the Dicer site. This reorganization occurs irrespective of which triple state is initially populated, consistent with binding to the dynamic A-bulge itself rather than a single structural endpoint. The functional importance of this flexibility was underscored by an engineered mutant that abolishes A-bulge shuffling and triple formation through uridine insertion; this rigid construct eliminated C1 binding regardless of approach pathway. These findings demonstrate that selectivity arises from the RNA’s capacity to sample productive binding geometries, rather than from simple intercalation or chemical complementarity alone, distinguishing RNA-selective ligands from promiscuous nucleic-acid binders.^61^

Thermodynamic analysis using Hamiltonian replica exchange metadynamics further supports this two-step recognition mechanism (**Figure 7**). *Wild type* pre-miR-377 exhibits a well-defined bound-state free energy minimum (**Figure 7A**), whereas the rigid mutant displays unfavorable binding energetics (ΔG ≈ 4 kcal·mol⁻¹ relative to unbound states). Importantly, ligand binding flattens the pre-miR-377 free energy landscape, lowering barriers between triple states and stabilizing partially disordered intermediates (**Figure 7B-C**). Optimal binding occurs when the RNA occupies conformations along the exchange pathway between triple states rather than when locked into either endpoint. This ensemble redistribution provides a mechanistic explanation for how C1 shifts the population away from Dicer-competent geometries, particularly those presenting the correct minor groove and 5′-phosphate architecture, thereby reducing miR-377 maturation and enabling VEGFA restoration in cells.

More broadly, these results suggest a generalizable paradigm for RNA-targeted small-molecule discovery: ligands can function by forcing RNAs into processing-incompatible folded states that are nonetheless energetically accessible within the native ensemble. Rather than inhibiting RNases through direct competition, small molecules can bias RNA folding landscapes toward geometries that fail quality-control checkpoints and are preferentially routed toward decay or inefficient processing. In this view, conformational frustration becomes a design principle. RNAs that rely on dynamic bulges, shuffling base pairs, or transient tertiary interactions for enzymatic recognition may be especially amenable to this strategy, whereas rigid, uniformly helical RNAs may be intrinsically resistant to small-molecule modulation.

In conclusion, we demonstrate that C1 targets pre-miR-377 through pathway-dependent engagement of a dynamic A-bulge, followed by ligand-induced redistribution of the conformational ensemble toward Dicer-incompatible states. The identification of conserved RNA triple motifs at this site, shared across phylogenetically distant regulatory RNAs, reveals fundamental architectural features that can be exploited for selective targeting. By integrating high-resolution NMR with experimentally refined ensemble modeling, this work provides both a mechanistic blueprint for pre-miRNA inhibition and a broader framework for identifying RNAs whose intrinsic dynamics render them tractable to small-molecule intervention.

## MATERIAL AND METHODS

### Sample design

In NMR, the limited variability of nucleobases results in poor spectral dispersion, complicating peak assignment and interpretation. Additionally, the dynamic nature of pre-miRs, particularly with flexible secondary structures such as loops or bulges, requires advanced time-consuming techniques. The overlap of ribose proton signals adds another layer of complexity, necessitating advanced NMR strategies and sample preparation to resolve these densely packed regions. The initial pre-miR-377 used was a 45-nucleotide containing an extra double GC pair resulting from enzymatic synthesis at the 5’ and 3’ ends. To optimize NMR data quality for MD incorporation, we strategically truncated pre-miR-377 45-nucleotide long sequence to 25 nucleotides (**Figure S17**), centered around the Dicer binding site. Additional GC pair was added at the 5’ and 3’ ends to ensure successful synthesis. The numbering was thus based on the 27-nucleotide long RNA, the first base in numbered 1. Our labelling strategy was guided by predicted secondary structures generated using mfold^62^ and RNAfold^63^. Leveraging typical chemical shift distributions from previous RNA NMR studies, we designed the labelling to gain insights into the base-pair pattern while avoiding overlapping signals.

### 5F-rFU labelled sample synthesis, purification and characterization

Reagents and phosphoramidites were purchased from Sigma-Aldrich part of the Merck group, TCI chemicals, WuXi AppTec, and used as received. The 5-fluoro-uridine phosphoramidites were synthesized in-house following standard procedure. Acetonitrile used for oligonucleotide synthesis was purchased as LiChrosolv grade from Sigma-Aldrich and further purified and dried by pushing through a MBRAUN SPS Compact using nitrogen gas immediately prior to use.

The ribo-fluorinated samples were synthesized on a MerMade48 instrument (Biosearch Technologies) at 5 µmol scale using resin functionalized with a universal linker (CUTAG CPG, Sigma Aldrich). Phosphoramidites were prepared as 0.1 M solutions in dry MeCN. Couplings were carried out using 0.25 M 5-[3,5-Bis(trifluoromethyl)phenyl]-1H-tetrazole (Activator 42®) in MeCN for 4×9 min. Oxidation was carried out with 0.02 M iodine in tetrahydrofuran/water/pyridine 90.54/9.05/0.41 (v/v/v) for 1 min. Unreacted 5’-hydroxyls were capped using a mixture of acetic anhydride / tetrahydrofuran 9.1/90.9 (v/v) (Cap A) and THF/NMI/pyridine 8/1/1 (v/v) (Cap B) for 1 min. Trityl groups were removed using 3% dichloroacetic acid (DCA) in toluene. After final detritylation, the 2-cyanoethyl protecting groups were removed using 20% diethylamine in MeCN. Following chain assembly, the solid-support-bound oligonucleotides were treated with a 1:1 mixture of 30% ammonium hydroxide and 40% aqueous methylamine at 65 °C for 60 min. Subsequently, 200 µL of 1 M Tris-HCl was added. The mixture was freeze-dried, and the resulting intermediate was treated with tetrabutylammonium fluoride in tetrahydrofuran for 24 h at room temperature. After deprotection, 200 µL of sodium acetate (pH 5.2) was added, and the oligonucleotides were precipitated with *tert*-butanol and ethanol, then dialyzed using a 10 kDa cut-off spin column.

The crude RNA was purified by reverse-phase high-performance liquid chromatography, Water LC 150, from Waters instrument (Milford), using a XBridge BEH C18 OBD Prpe Column, 130A 5um, 10 × 250 mm as column and ion pairing chromatography (Triethylammonium acetate buffer, 0.1M, pH 7.0). Purification was performed using a linear gradient from 5 to 20% acetonitrile:100mM TEAA (aq) over 30 min.

Identity and purity of oligonucleotides were confirmed by liquid chromatography-mass spectrometry (LC-MS) using an Acquity I-class LC system, equipped with a PDA and coupled to an RDa via heated electrospray (BioAccord) by the Waters Corporation. Yields were quantified by nanodrop.

### Sample Preparation

Pre-miR-377 samples for NMR experiments were prepared in 3mm or Shigemi NMR sample tubes. The samples consisted of selectively labelled, unlabeled, and fully labelled 27-nucleotide long pre-miR-377, dissolved in either 100% D_2_O or 90% H_2_O/10% D_2_O. The buffer conditions were set at 25 mM K-phosphate at pH 6.2, with additional 50 mM KCl and 5 mM MgCl2. Sample concentrations ranged from 400 μM to 2.2 mM. For 1D binding experiments, the stock solution of unlabeled pre-miR-377 was prepared at a concentration of 150 µM. The compound C1 was diluted in 20 mM sodium phosphate (NaPi) at pH 7.4, 150 mM NaCl, and 10% D_2_O. For 2D binding experiments, buffer was identical to the one used for structural investigation.

### NMR Spectroscopy

NMR experiments were conducted on Bruker spectrometers of 600 and 800 MHz frequencies, equipped with 5-mm cryogenic probes, located at the AstraZeneca NMR Centre of Excellence in Gothenburg and the Swedish NMR Centre of the University of Gothenburg. Spectra were collected at 283 K and 298K. For structural analysis, a series of ^1^H, ^15^N and ^1^H, ^13^C-SOFAST HMQC were recorded on selectively labelled, unlabeled, and fully labelled samples. Assignment of non-exchangeable protons was facilitated through 2D TOCSY acquisition for all samples with a mixing time of 10, 35 and 100 ms, and base-pairing pattern was identified using HNN-COSY (U-A and G-C) and selective HNN-COSY (A-U). NOESY experiments were performed in both H_2_O and D_2_O environments with various mixing times including 50 ms, 100 ms, and 150 ms. Detailed NMR experimental parameters, such as spectral widths, time domain points, and acquisition parameters for each experiment type, can be found in Table S4.

For binding experiments, a series of 10 X 1D proton Carr-Purcell-Meiboom-Gill (CPMG) experiments were performed at 298K by sequentially adding 1 µL aliquots of the pre-miR-377 stock solution to a sample containing compound C1. The total reaction volume was 160 ul. Additional 5 x 2D [1H,13C]-SOFAST HMQC experiments were conducted at 283K on a [15N, 13C]-uniformly labelled 50 µM pre-miR-377 sample, with varying concentrations of compound C1: 0, 25, 50, 75, and 100 µM.

For F-functionalized samples, 19F NMR experiments were acquired with and without proton decoupling. Experiments were performed at 298K on different F-labelled pre-miR-377 samples, with a concentration of 200 µM RNA.

### Data Processing and Analysis

NMR data was processed using NMRPipe or Topspin and analyzed using CCPNMR. Proton chemical shifts were internally referenced to Sodium trimethylsilylpropanesulfonate (DSS), while carbon chemical shifts were indirectly referenced relative to the proton shifts.

### Structural modeling and MD simulations

An initial ensemble of 750 conformers of pre-miR-377 was generated using the fragment assembly and all-atom refinement protocol FARFAR2,^64^ starting from the most populated secondary structure modelled by NMR, and enforcing base pairs in the stem. The top-scoring pre-miR-377 model was selected as the seed for independent all-atom molecular dynamics (MD) simulations. The same approach has been used to generate 50 conformers of pre-miR-377 mutant (Uridine insertion between A24 and U25). Following solvation in a dodecahedral box with 1.0 nm buffer, K⁺ and Mg²⁺ ions were added to neutralize the system and reproduce the experimental NMR buffer (25 mM potassium phosphate, pH 6.2, 50 mM KCl, 5 mM MgCl₂). Two combinations of force fields and solvent models were tested: (i) Amber χOL3 nucleic-acid correction^46–49^ with OPC water,^65^ and (ii) the DESRES RNA force field^50^ with TIP4P water model.^66,67^ Monovalent ions were modeled with Joung-Cheatham parameters,^68^ and divalent cations with the Li-Merz 12-6 parameters.^69^

System preparation followed a standardized equilibration protocol: (i) steepest-descent energy minimization of solvent and ions with positional restraints on RNA heavy atoms; (ii) 1.5 ns NVT equilibration at 298 K with 500 kJ mol⁻¹ nm⁻² restraints (Δt = 1 fs); (iii) 2.5 ns NPT equilibration at 1 bar with the same restraints (Δt = 2 fs); (iv) five sequential 1.25 ns NPT stages with progressively reduced restraints; and (v) a final 5 ns NPT run without restraints. Velocity-rescaling thermostat^70^ with a relaxation time τ= 0.1 ps, and Parrinello-Rahman barostat^71^ at reference pressure of 1 atm with τ= 2 ps were used.

To enhance sampling of conformational space, production simulations employed Hamiltonian replica exchange MD (H-REMD)^72,73^ combined with well-tempered metadynamics,^74^ implemented via PLUMED 2.9^75^ patched into GROMACS 2021.4.^76^ For each system, 12 replicas were distributed across a Hamiltonian ladder by scaling nonbonded interaction terms, spanning an effective temperature range of 310-460 K. Exchange attempts were attempted every 300 ps with a Metropolis criterion, yielding ∼14-40% acceptance (**Figure S8A**). The radius of gyration of the RNA was used as collective variable, with gaussians’ height 0.5 kJ mol^−1^, σ of 0.06 nm, deposition rate of 500 steps and bias factor of 5.

The exact same protocol was used to perform three independent H-REMD simulations in the presence of the small molecule C1 and: i) pre-miR-377 centroid of cluster #1; ii) pre-miR-377 centroid of cluster #6; iii) pre-miR-377 mutant top scoring FARFAR model. For these systems, both the RNA radius of gyration (gaussians’ height 2.5 kJ mol^−1^, σ of 0.06 nm) and the distance between the center of mass of C1 and the Dicer Site (A3, A4, U5, A24, U25; gaussians’ height 2.5 kJ mol^−1^, σ of 0.03 nm) were used as collective variables, with deposition rate of 500 steps, and bias factor of 18. The General AMBER Force Field (GAFF)^77^ was used to model bonded and nonbonded interactions of C1. The protonation state of C1 was assigned based on the benzimidazole pKa, determined both computationally (Jaguar, default QM scheme) and experimentally by ionization capillary electrophoresis, and considering the NMR buffer pH of 6.5.

To monitor conformational transitions of the Dicer site in the presence of C1, we defined a path CV over nucleotides G2-U5 and A23-C26. The path endpoints were the two reference structures sampled with MD: the A3-U25•A4 triple (cluster #1; Figure 4B) and the G2-C26•A3 triple (cluster #6; Figure 4C). These endpoints are connected by 13 intermediate states uniformly spaced in eRMSD, ensuring a constant increment of 0.10 eRMSD per path node (total path length 1.53 eRMSD).

### Steered MD simulations

A set of 200 independent steered MD simulations were performed for each of the following systems: pre-miR-377 cluster #1 centroid, pre-miR-377 cluster #6 centroid, and pre-miR-377 mutant top FARFAR model with C1 positioned 12Å from the either the minor or the major groove, resulting in a total of six different systems. Here, only the force field combination fitting better the NMR was used: Amber χOL3 nucleic-acid correction^46–49^ with OPC water^65^, while monovalent ions were modeled with Joung-Cheatham parameters,^68^ and divalent cations with the Li-Merz 12-6 parameters.^69^

Each simulation extended for 5 ns, yielding a total simulation time of 6 μs across all systems. The simulations employed a three-step biasing protocol designed to facilitate RNA-ligand recognition under controlled conditions. During the initial 2.5 ns, a moving distance restraint of 500 kJ·mol^-1^·Å^-1^ was applied between the center of mass of C1 and the Dicer cleavage site, targeting zero distance. Simultaneously, the RNA conformation was restrained over the input coordinates for each system using positional restraints with a force constant of 500 kJ·mol⁻¹·Å^-1^. The C1-Dicer site distance restraint was removed at 2.5 ns, while RNA conformational restraints remained active until 3.5 ns to allow initial ligand accommodation. Following this transition period, all restraints were eliminated, enabling 1.5 ns of unbiased equilibrium sampling to capture RNA-small molecule recognition events under physiologically relevant conditions. Final coordinates from each simulation were collected to generate ensemble datasets of 200 structures per system, which served as the basis for subsequent structural analyses. Trajectory analysis was performed with BARNABA^78^ and PLUMED.^75^

### Ensemble Reweighting with BME

All ensembles generated with FARFAR2 and MD simulations were refined against experimental NMR data using the Bayesian/Maximum Entropy reweighting approach implemented in the BME python library.^40^ In this framework, MD-derived ensembles represent the prior distribution, while NMR observables define the likelihood. Specifically, interproton distances from NOESY spectra (averaging seven restraints per nucleotide, 175 non-redundant restraints in total) were used as reweighting targets. For each trajectory frame, back-calculated distances were compared to experimental values. The BME algorithm optimizes conformer weights *(w)* by minimizing:

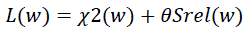

Where *χ*2(*w*) measures deviations between experimental observables and reweighted ensemble averages, while *Srel*(*w*) is the relative Shannon entropy penalizing deviations from the prior. The balance between fitting and prior preservation is controlled by *θ* chosen after cross-validation. Upon inspection of cross-validated ensembles, a unique value of *θ* = 1.13 was applied across all reweighting runs to avoid ensemble-specific overfitting (**Figure S10**).

## AUTHOR CONTRIBUTION

G.L.S., C.N.C., A.H., and S.G. conceived the research. J.M., E.M. and C.N.C. designed experiments and collected and analyzed the data. J.M., G.L.S., E.M., and C.N.C. interpreted the data. D.D., M.B., and R.Q., synthetized the RNA constructs. G.L.S. and C.N.C. provided supervision. A.I.F provided technical support. J.M. and C.C. wrote the original draft. All authors contributed to the drafting of the manuscript and approved the final version.

## ACKNOWLEDGMENT

J.M., E.M., and D.D. are fellows of the AstraZeneca R&D postdoc program. This work was supported by the National Institutes of Health (R35NS116846 to M.D.D.)

## DATA AVAILABILITY

The data that support the findings of this study are available from the corresponding authors upon reasonable request.

## REFERENCES

1. Włodarczyk, M. et al. The role of miRNAs in the pathogenesis, diagnosis, and treatment of colorectal cancer and colitis-associated cancer. Clin. Exp. Med. 25, 1–19 (2025).

2. Hashimoto, N. & Tanaka, T. Role of miRNAs in the pathogenesis and susceptibility of diabetes mellitus. J. Hum. Genet. 62, 141–150 (2017).

3. Feliciano, A., Sánchez-Sendra, B., Kondoh, H. & Lleonart, M. E. MicroRNAs regulate key effector pathways of senescence. J. Aging Res. 2011, (2011).

4. Lin, S. & Gregory, R. I. MicroRNA biogenesis pathways in cancer. Nat. Rev. Cancer 15, 321–333 (2015).

5. Sun, L. L., Li, W. D., Lei, F. R. & Li, X. Q. The regulatory role of microRNAs in angiogenesis-related diseases. J. Cell. Mol. Med. 22, 4568–4587 (2018).

6. Zeng, Y. Principles of micro-RNA production and maturation. Oncogene 25, 6156–6162 (2006).

7. Childs-Disney, J. L. et al. Targeting RNA structures with small molecules. Nat. Rev. Drug Discov. 21, 736–762 (2022).

8. Warner, K. D., Hajdin, C. E. & Weeks, K. M. Principles for targeting RNA with drug-like small molecules. Nat. Rev. Drug Discov. 17, 547–558 (2018).

9. Disney, M. D. Targeting RNA with Small Molecules To Capture Opportunities at the Intersection of Chemistry, Biology, and Medicine. J. Am. Chem. Soc. 141, 6776–6790 (2019).

10. Velagapudi, S. P. et al. Design of a small molecule against an oncogenic noncoding RNA. Proc. Natl. Acad. Sci. U. S. A. 113, 5898–5903 (2016).

11. Velagapudi, S. P., Gallo, S. M. & Disney, M. D. Sequence-based design of bioactive small molecules that target precursor microRNAs. Nat. Chem. Biol. 10, 291–297 (2014).

12. Costales, M. G. et al. A Designed Small Molecule Inhibitor of a Non-Coding RNA Sensitizes HER2 Negative Cancers to Herceptin. J. Am. Chem. Soc. 141, 2960–2974 (2019).

13. Staedel, C. et al. Modulation of oncogenic miRNA biogenesis using functionalized polyamines. Sci. Rep. 8, 1–13 (2018).

14. Becquart, C. et al. Exploring Heterocycle-Spermine Conjugates as Modulators of Oncogenic microRNAs Biogenesis. ACS Omega 3, 16500–16508 (2018).

15. Maucort, C. et al. Design and Implementation of Synthetic RNA Binders for the Inhibition of miR-21 Biogenesis. ACS Med. Chem. Lett. (2021) doi:10.1021/acsmedchemlett.0c00682.

16. Zhang, P. et al. Reprogramming of Protein-Targeted Small-Molecule Medicines to RNA by Ribonuclease Recruitment. J. Am. Chem. Soc. 143, 13044–13055 (2021).

17. Ratni, H. et al. Discovery of Risdiplam, a Selective Survival of Motor Neuron-2 (SMN2) Gene Splicing Modifier for the Treatment of Spinal Muscular Atrophy (SMA). J. Med. Chem. 61, 6501–6517 (2018).

18. Sivaramakrishnan, M. et al. Binding to SMN2 pre-mRNA-protein complex elicits specificity for small molecule splicing modifiers. Nat. Commun. 8, (2017).

19. Bottaro, S. & Lindorff-Larsen, K. Biophysical experiments and biomolecular simulations: A perfect match? Science vol. 361 355–360 at 10.1126/science.aat4010 (2018).

20. Salmon, L., Yang, S. & Al-Hashimi, H. M. Advances in the Determination of Nucleic Acid Conformational Ensembles. Annu. Rev. Phys. Chem. 65, 293–316 (2014).

21. Al-Hashimi, H. M. & Walter, N. G. RNA dynamics: it is about time. Curr. Opin. Struct. Biol. 18, 321–329 (2008).

22. Dethoff, E. A., Chugh, J., Mustoe, A. M. & Al-Hashimi, H. M. Functional complexity and regulation through RNA dynamics. Nature 482, 322–330 (2012).

23. Cruz, J. A. et al. RNA-Puzzles: A CASP-like evaluation of RNA three-dimensional structure prediction. Rna 18, 610–625 (2012).

24. Silvestri, I. et al. Targeting the conserved active site of splicing machines with specific and selective small molecule modulators. Nat. Commun. 15, 4980 (2024).

25. Manigrasso, J. et al. Visualizing group II intron dynamics between the first and second steps of splicing. Nat. Commun. 11, (2020).

26. Martino, G., Manigrasso, J., La Sala, G., Marcia, M. & De Vivo, M. Controlled dynamic remodeling of the spliceosome active site enables the first step of splicing. Proc. Natl. Acad. Sci. U. S. A. 123, 1–12 (2026).

27. Jadhav, S. et al. Dynamic assembly of a large multidomain ribozyme visualized by cryo-electron microscopy. Nat. Commun. 16, 10195 (2025).

28. Mertinkus, K. R. et al. Dissecting the Conformational Heterogeneity of Stem-Loop Substructures of the Fifth Element in the 5′-Untranslated Region of SARS-CoV-2. J. Am. Chem. Soc. 146, 30139–30154 (2024).

29. Hawkes, E. J. et al. COOLAIR Antisense RNAs Form Evolutionarily Conserved Elaborate Secondary Structures. Cell Rep. 16, 3087–3096 (2016).

30. Stelzer, A. C. et al. Discovery of selective bioactive small molecules by targeting an RNA dynamic ensemble. Nat. Chem. Biol. 7, 553–559 (2011).

31. Oxenfarth, A. et al. Integrated NMR/Molecular Dynamics Determination of the Ensemble Conformation of a Thermodynamically Stable CUUG RNA Tetraloop. J. Am. Chem. Soc. 145, 16557–16572 (2023).

32. Sponer, J. et al. RNA structural dynamics as captured by molecular simulations: A comprehensive overview. Chem. Rev. 118, 4177–4338 (2018).

33. Marušič, M., Toplishek, M. & Plavec, J. NMR of RNA - Structure and interactions. Curr. Opin. Struct. Biol. 79, (2023).

34. Marcia, M., Manigrasso, J. & De Vivo, M. Finding the Ion in the RNA-Stack: Can Computational Models Accurately Predict Key Functional Elements in Large Macromolecular Complexes? J. Chem. Inf. Model. 61, 2511–2515 (2021).

35. Muscat, S., Martino, G., Manigrasso, J., Marcia, M. & De Vivo, M. On the Power and Challenges of Atomistic Molecular Dynamics to Investigate RNA Molecules. J. Chem. Theory Comput. (2024) doi:10.1021/acs.jctc.4c00773.

36. Manigrasso, J., Marcia, M. & De Vivo, M. Computer-aided design of RNA-targeted small molecules: A growing need in drug discovery. Chem 1–24 (2021) doi:10.1016/j.chempr.2021.05.021.

37. Connelly, C. M., Moon, M. H. & Schneekloth, J. S. The Emerging Role of RNA as a Therapeutic Target for Small Molecules. Cell Chem. Biol. 23, 1077–1090 (2016).

38. Nagasawa, R. et al. Crystallographic analysis of G-clamp–RNA complex assisted by large scale RNA-binding profile. Chem. Commun. 61, 1120–1123 (2025).

39. Haniff, H. S. et al. Design of a small molecule that stimulates vascular endothelial growth factor A enabled by screening RNA fold–small molecule interactions. Nat. Chem. 12, 952–961 (2020).

40. Bottaro, S., Bengtsen, T. & Lindorff-Larsen, K. Integrating Molecular Simulation and Experimental Data: A Bayesian/Maximum Entropy Reweighting Approach. in In Structural Bioinformatics: Methods and Protocols, Methods in Molecular Biology 219–240 (Humana Press Inc., 2020). doi:10.1007/978-1-0716-0270-6_15.

41. Hennig, M., Scott, L. G., Sperling, E., Bermel, W. & Williamson, J. R. Synthesis of 5-fluoropyrimidine nucleotides as sensitive NMR probes of RNA structure. J. Am. Chem. Soc. 129, 14911–14921 (2007).

42. Zhao, S. et al. Dynamics of base pairs with low stability in RNA by solid-state nuclear magnetic resonance exchange spectroscopy. iScience 25, (2022).

43. Cavanagh, J., Skelton, N. J., Fairbrother, W. J., Rance, M. & Palmer, A. G. Protein NMR Spectroscopy Principles and Practice. Acad. Press (2006).

44. Ryabov, Y., Schwieters, C. D. & Clore, G. M. Determination with Sparse Distance Restraints. J. Am. Chem. Soc. 0–3 (2011).

45. Kneller, J. M., Lu, M. & Bracken, C. An effective method for the discrimination of motional anisotropy and chemical exchange. J. Am. Chem. Soc. 124, 1852–1853 (2002).

46. Pérez, A. et al. Refinement of the AMBER force field for nucleic acids: Improving the description of α/γ conformers. Biophys. J. 92, 3817–3829 (2007).

47. Zgarbová, M. et al. Refinement of the Cornell et al. Nucleic acids force field based on reference quantum chemical calculations of glycosidic torsion profiles. J. Chem. Theory Comput. 7, 2886–2902 (2011).

48. Steinbrecher, T., Latzer, J. & Case, D. A. Revised AMBER parameters for bioorganic phosphates. J. Chem. Theory Comput. 8, 4405–4412 (2012).

49. Bergonzo, C. & Cheatham, T. E. Improved Force Field Parameters Lead to a Better Description of RNA Structure. J. Chem. Theory Comput. 11, 3969–3972 (2015).

50. Tan, D., Piana, S., Dirks, R. M. & Shaw, D. E. RNA force field with accuracy comparable to state-of-the-art protein force fields. Proc. Natl. Acad. Sci. U. S. A. 115, E1346–E1355 (2018).

51. Bottaro, S., Bussi, G. & Lindorff-Larsen, K. Conformational Ensembles of Noncoding Elements in the SARS-CoV-2 Genome from Molecular Dynamics Simulations. J. Am. Chem. Soc. 143, 8333–8343 (2021).

52. Bernetti, M., Hall, K. B. & Bussi, G. Reweighting of molecular simulations with explicit-solvent SAXS restraints elucidates ion-dependent RNA ensembles. Nucleic Acids Res. 49, (2021).

53. Geng, A. et al. An RNA excited conformational state at atomic resolution. Nat. Commun. 14, (2023).

54. Westhof, E., Sun, H., Bu, F. & Miao, Z. The RNA -Puzzles Assessments of RNA - Only Targets in CASP16. Proteins Struct. Funct. Bioinforma. 1–12 (2025) doi:10.1002/prot.70052.

55. Palmer, A. G. & Koss, H. Chemical Exchange. Methods in Enzymology vol. 615 (Elsevier Inc., 2019).

56. Abu Almakarem, A. S., Petrov, A. I., Stombaugh, J., Zirbel, C. L. & Leontis, N. B. Comprehensive survey and geometric classification of base triples in RNA structures. Nucleic Acids Res. 40, 1407–1423 (2012).

57. Slaymaker, I. M. et al. High-Resolution Structure of Cas13b and Biochemical Characterization of RNA Targeting and Cleavage. Cell Rep. 26, 3741–3751.e5 (2019).

58. Manigrasso, J., De Vivo, M. & Palermo, G. Controlled Trafficking of Multiple and Diverse Cations Prompts Nucleic Acid Hydrolysis. ACS Catal. 11, 8786–8797 (2021).

59. Ahsan, M., Pindi, C. & Palermo, G. Emerging Mechanisms of Metal-Catalyzed RNA and DNA Modifications. Annu. Rev. Phys. Chem. 76, 497–518 (2025).

60. Narayanaswamy, N. et al. Sequence-specific recognition of DNA minor groove by an NIR-fluorescence switch-on probe and its potential applications. Nucleic Acids Res. 43, 8651–8663 (2015).

61. Dervan, P. B. Molecular recognition of DNA by small molecules. Bioorganic Med. Chem. 9, 2215–2235 (2001).

62. Zuker, M. Mfold web server for nucleic acid folding and hybridization prediction. Nucleic Acids Res. 31, 3406–3415 (2003).

63. Gruber, A. R., Lorenz, R., Bernhart, S. H., Neuböck, R. & Hofacker, I. L. The Vienna RNA websuite. Nucleic Acids Res. 36, 70–74 (2008).

64. Watkins, A. M., Rangan, R. & Das, R. FARFAR2: Improved De Novo Rosetta Prediction of Complex Global RNA Folds. Structure 28, 963–976.e6 (2020).

65. Izadi, S., Anandakrishnan, R. & Onufriev, A. V. Building Water Models: A Different Approach. J. Phys. Chem. Lett. 5, 3863–3871 (2014).

66. Jorgensen, W. L., Chandrasekhar, J., Madura, J. D., Impey, R. W. & Klein, M. L. Comparison of simple potential functions for simulating liquid water. J. Chem. Phys. 79, 926–935 (1983).

67. Jorgensen, W. L. & Madura, J. D. Temperature and size dependence for Monte Carlo simulations of TIP4P water. Mol. Phys. 56, 1381–1392 (1985).

68. Joung, I. S. & Cheatham, T. E. Molecular Dynamics Simulations of the Dynamic and Energetic Properties of Alkali and Halide Ions Using Water-Model-Specific Ion Parameters. J. Phys. Chem. B 113, 13279–13290 (2009).

69. Li, P., Roberts, B. P., Chakravorty, D. K. & Merz, K. M. J. Rational Design of Particle Mesh Ewald Compatible Lennard-Jones Parameters for +2 Metal Cations in Explicit Solvent. J. Chem. Theory Comput. 9, 2733–2748 (2013).

70. Bussi, G., Donadio, D. & Parrinello, M. Canonical sampling through velocity rescaling. J. Chem. Phys. 126, 14101 (2007).

71. Parrinello, M. & Rahman, A. Polymorphic transitions in single crystals: A new molecular dynamics method. J. Appl. Phys. 52, 7182–7190 (1981).

72. Sugita, Y. & Okamoto, Y. Replica-exchange multicanonical algorithm and multicanonical replica-exchange method for simulating systems with rough energy landscape. Chem. Phys. Lett. 329, 261–270 (2000).

73. Bussi, G. Hamiltonian replica exchange in GROMACS: a flexible implementation. Mol. Phys. 112, 379–384 (2014).

74. Barducci, A., Bussi, G. & Parrinello, M. Well-Tempered Metadynamics: A Smoothly Converging and Tunable Free-Energy Method. Phys. Rev. Lett. 100, 020603 (2008).

75. The PLUMED Consortium. Promoting transparency and reproducibility in enhanced molecular simulations. Nat. Methods 16, 670–673 (2019).

76. Abraham, M. J. et al. GROMACS: High performance molecular simulations through multi-level parallelism from laptops to supercomputers. SoftwareX 1–2, 19–25 (2015).

77. Wang, J., Wolf, R. M., Caldwell, J. W., Kollman, P. A. & Case, D. A. Development and testing of a general amber force field. J. Comput. Chem. 25, 1157–1174 (2004).

78. Bottaro, S. et al. Barnaba: Software for Analysis of Nucleic Acids Structures and Trajectories. RNA 25, 219–231 (2019).

