## supplemental material for "Conformational dynamics at the pre-miR-377 Dicer site governs selective small-molecule recognition"

### Conformational dynamics at the Dicer site enables selective small molecule recognition of microRNA-377 precursor

**This PDF file includes:**

Figures S1 to S21
Table S1 to S4
Synthesis of phosphoramidite

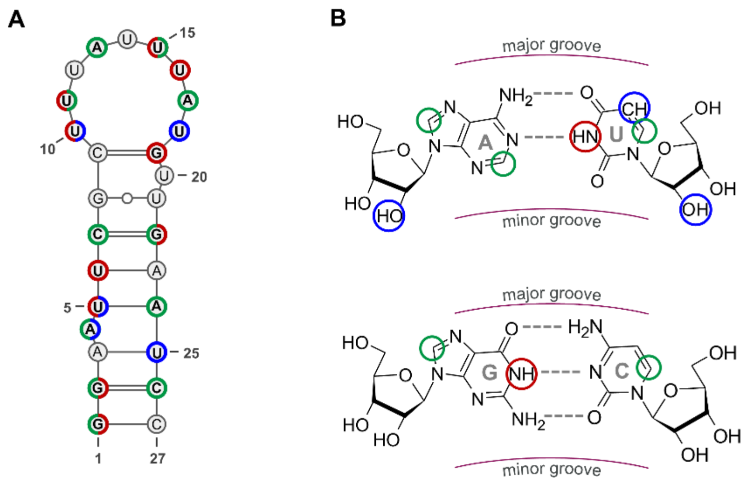

**Figure S1. Labeling scheme and nucleotide annotation used for pre-miR-377. (A)** Representative 2D secondary structure of pre-miR-377 construct used for NMR experiments indicating the positions of selectively introduced labels. Colored circles indicate $^{15}$N-H labeling (NH, red), $^{13}$C-H labeling (CH, green), $^{19}$F replacement of hydrogen (blue). The summary of all selectively labeled constructs used in this study is reported in Table 1. **(B)** Chemical structures highlighting the corresponding positions of the labels on AU and GC base pairs following same coloring scheme as panel A. Major- and minor-groove orientations are indicated for reference.

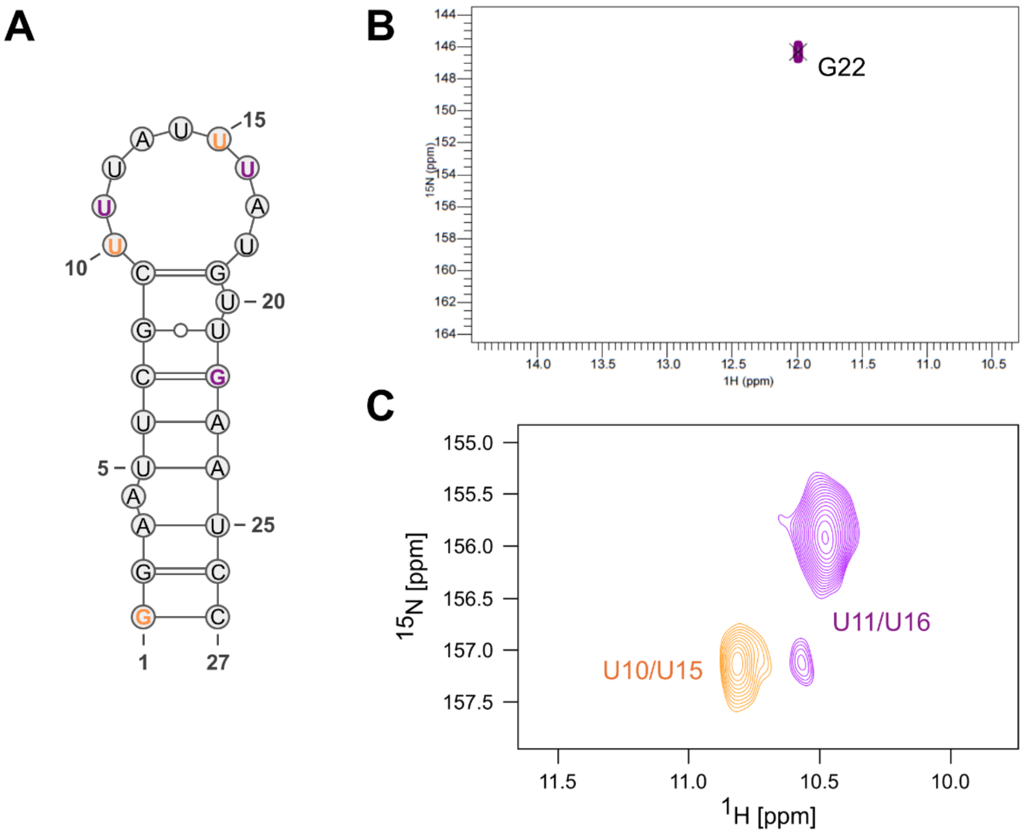

**Figure S2: Analysis of pre-miR-377 base-pairing network through selectively labelled nucleotides, focusing on the apical loop region. (A)** Secondary structure of pre-miR-377 highlighting selectively labelled nucleotides in two distinct colours, each corresponding to a unique sample and indicating the specific positions of the $^{15}N$ labels. **(B-C)** [$^{1}H$, $^{15}N$]-SOFAST-HMQC spectra at 283K for each sample, illustrating the base-pairing network and emphasizing the absence of strong interactions within the loop region.

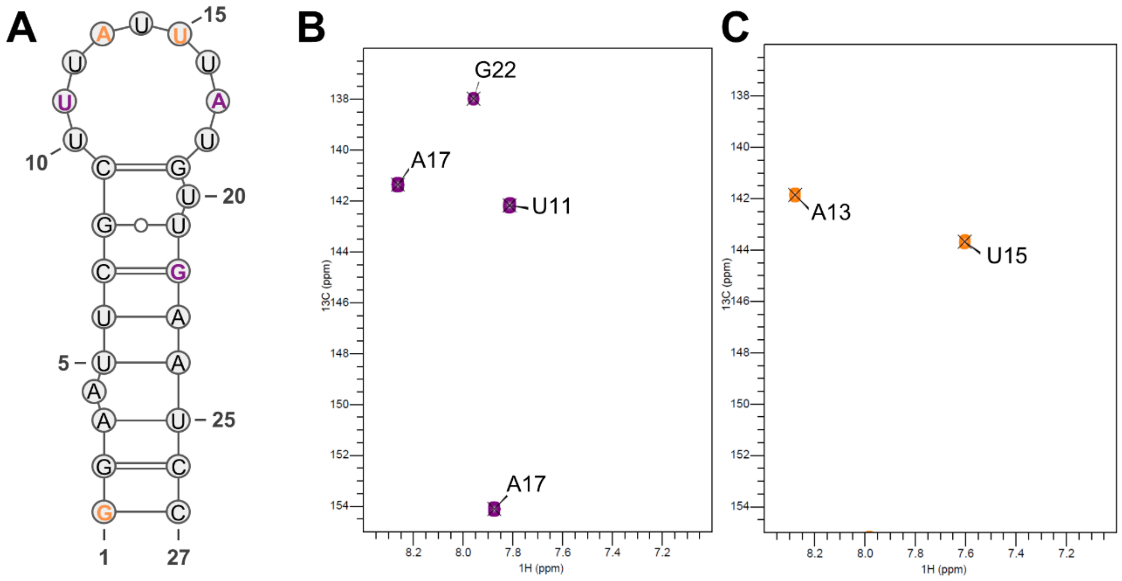

**Figure S3: Identification of pre-miR-377 resonances through selectively labelled nucleotides. (A)** Secondary structure of pre-miR-377, showcasing selectively labelled nucleotides in two distinct colours, each corresponding to a unique sample and indicating the specific positions of the $^{13}$C labels. **(B-C)** [$^{1}$H, $^{13}$C]-SOFAST-HMQC spectra at 283K for each sample. Both A13 and A17 H8 arise in the de-shielded region, aligning with previous observations that suggest their positioning within a loop region.

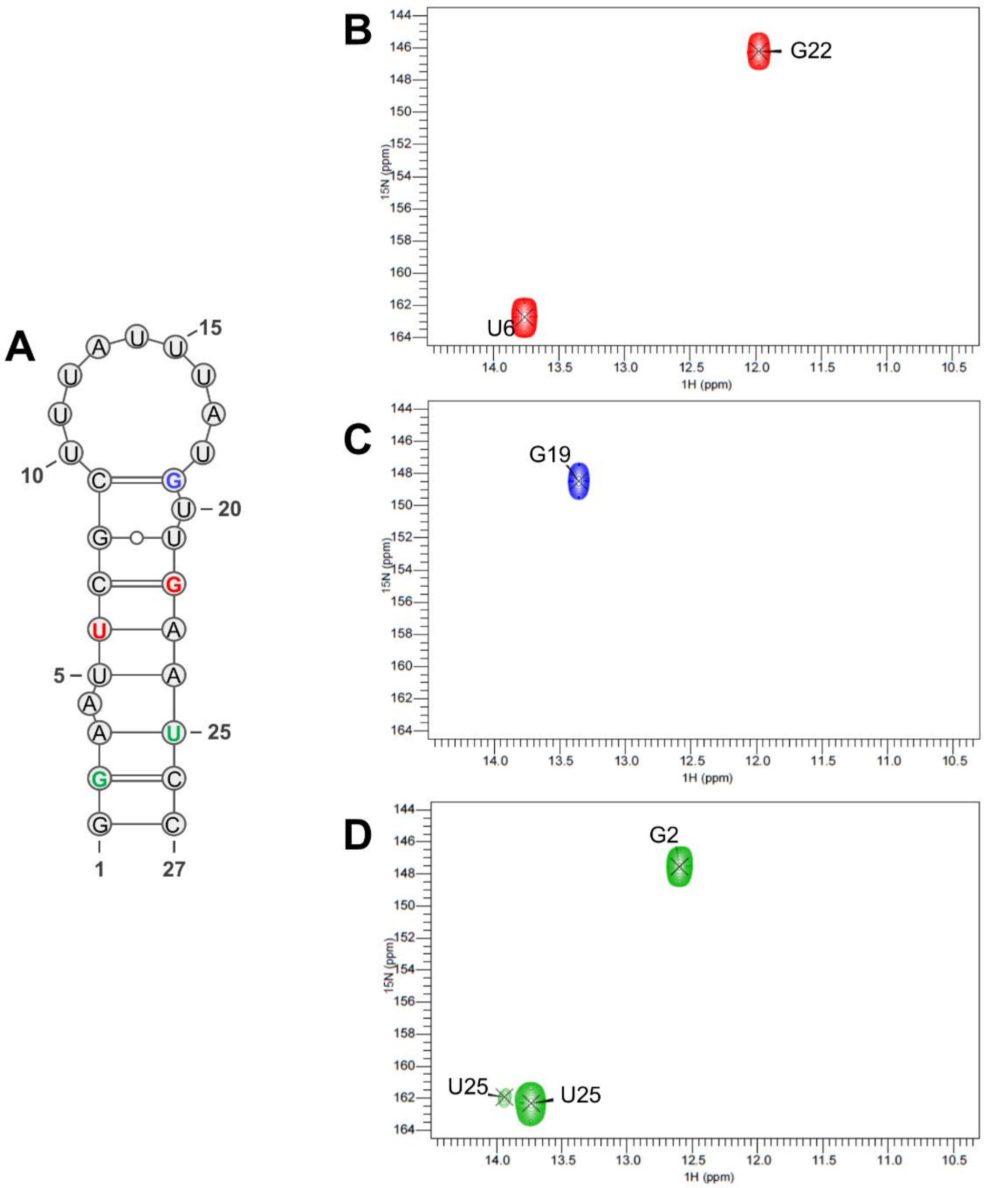

**Figure S4: Mapping of pre-miR-377 base-pairing network through selectively labelled nucleotides. (A)** Secondary structure of pre-miR-377, showcasing selectively labelled nucleotides in three distinct colours, each corresponding to a unique sample and indicating the specific positions of the $^{15}$N labels. **(B-D)** [$^{1}$H, $^{15}$N]-SOFAST-HMQC spectra at 283K for each sample, illustrating the base-pairing network within the pre-miR-377.

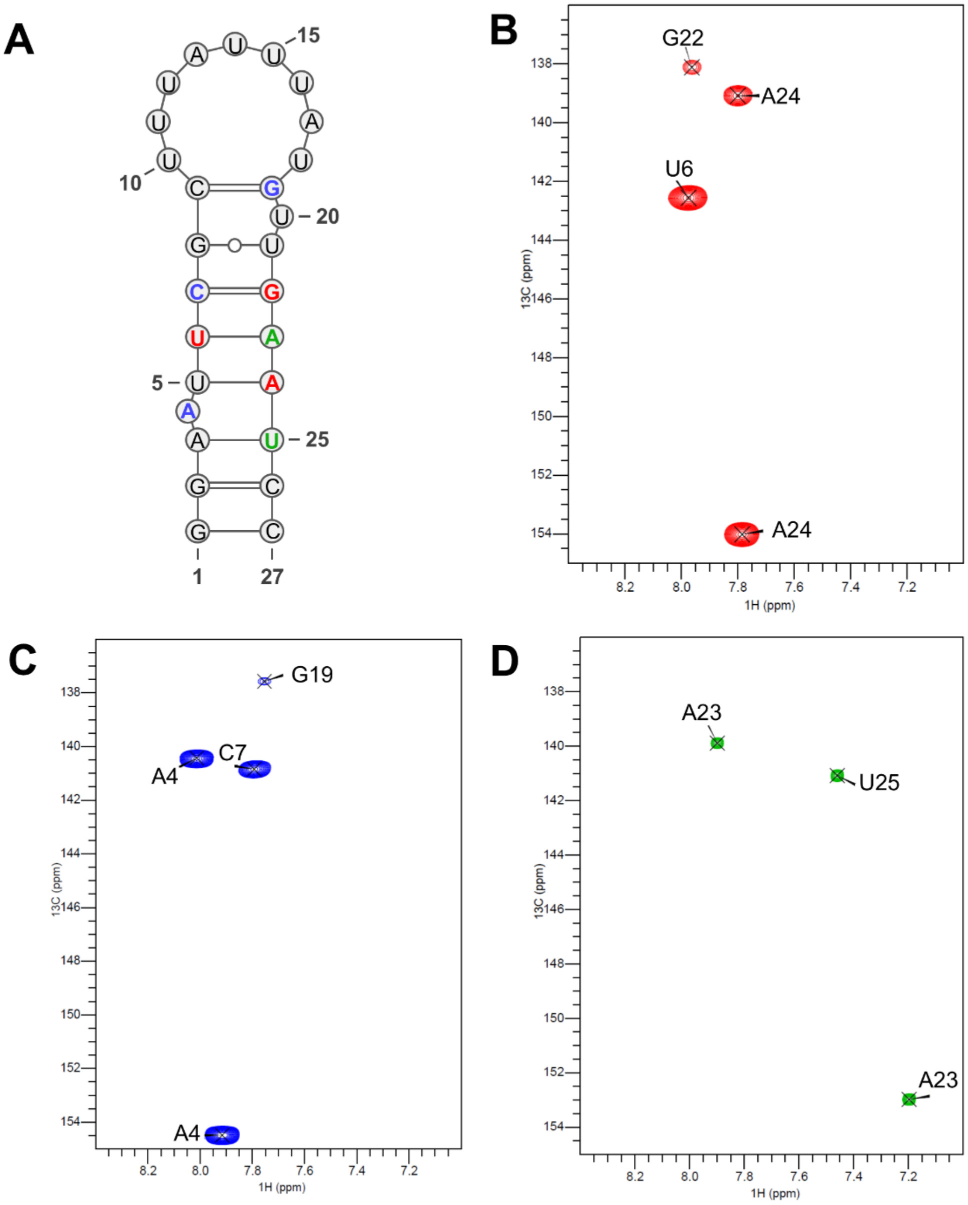

**Figure S5: Identification of pre-miR-377 resonances through selectively labelled nucleotides. (A)** Secondary structure of pre-miR-377, showcasing selectively labelled nucleotides in three distinct colours, each corresponding to a unique sample and indicating the specific positions of the $^{13}C$ labels. **(B-D)** [$^{1}H$, $^{13}C$]-SOFAST-HMQC spectra at 283K for each sample.

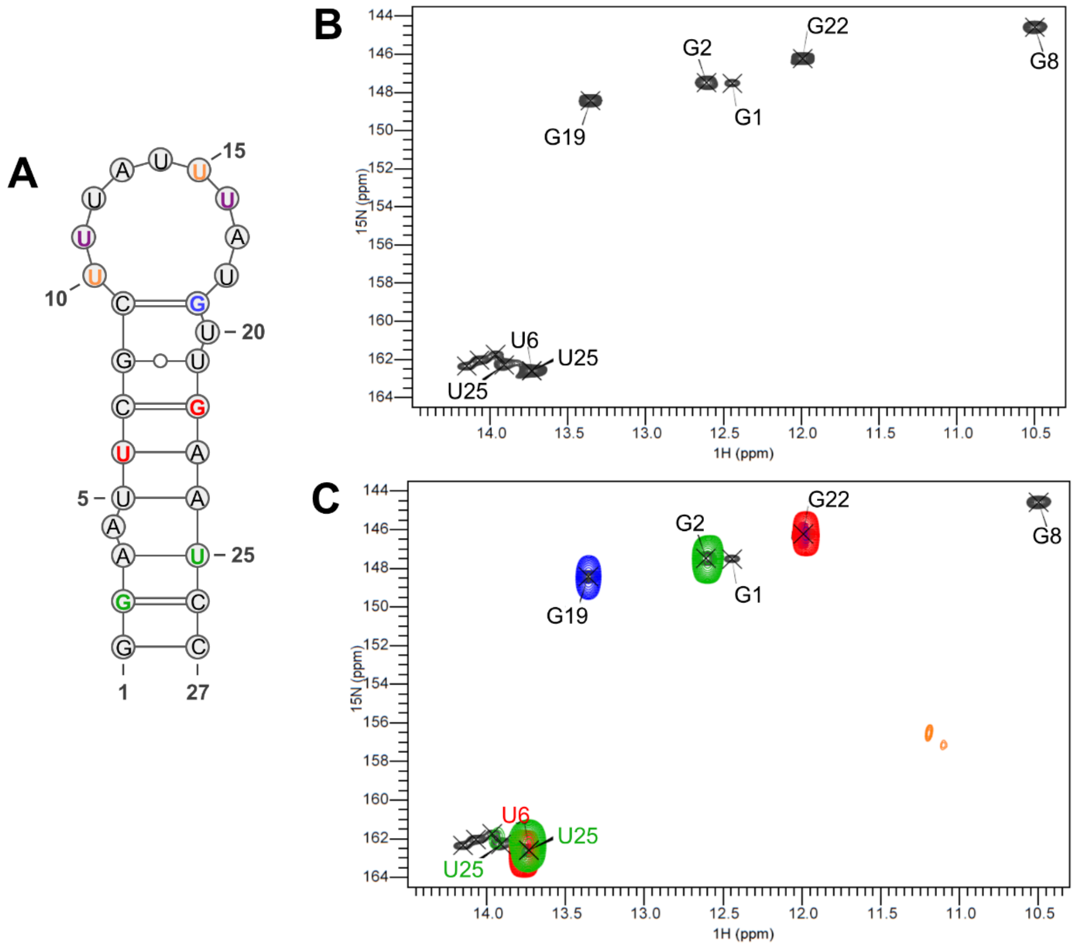

**Figure S6: Analysis of U-A interactions and their conformational states in pre-miR-377 structure. (A)** Secondary structure highlighting selectively labelled nucleotides in five distinct colours, each representing a different sample and indicating the specific positions of the $^{15}N$ labels. **(B)** [$^{1}H$, $^{15}N$]-SOFAST HMQC spectrum of the fully labelled sample, illustrating numerous U-A base pairs. **(C)** Superimposition of the fully labelled spectra and all selectively labelled spectra, demonstrating little overlap in the crowded U-A region. This indicates that the observed signals do not arise only from the loop but also from the Dicer site region exhibiting multiple conformers.

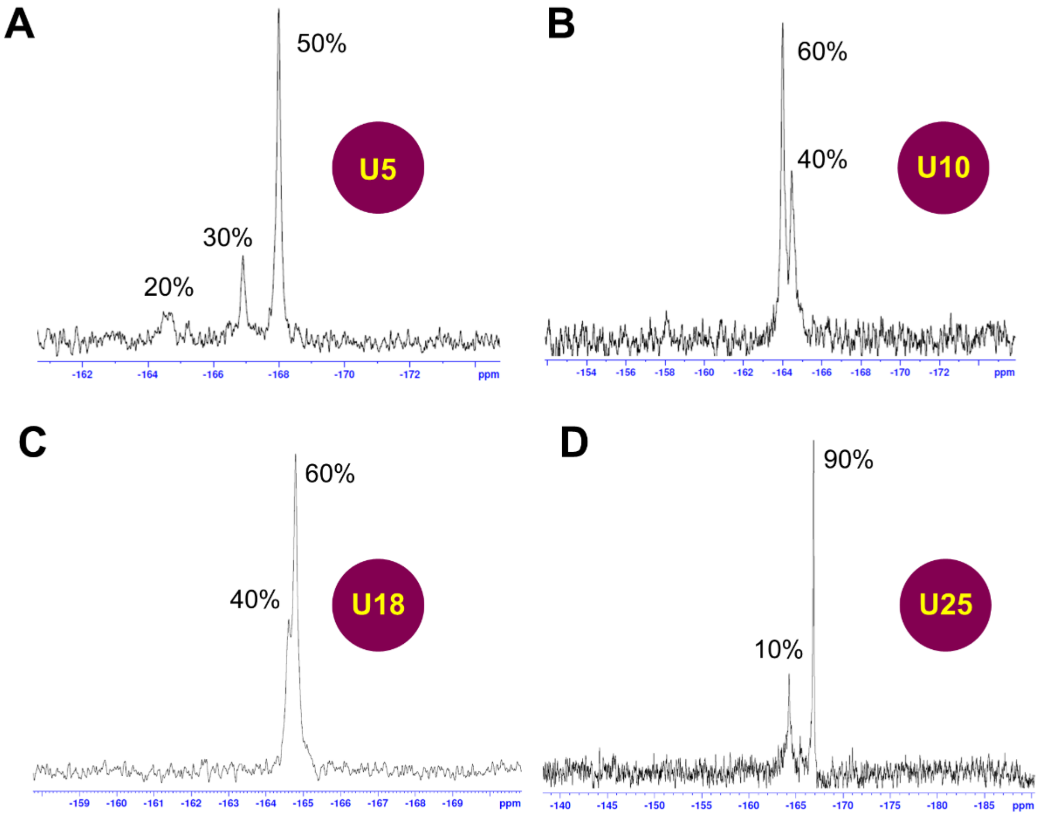

**Figure S7. Site-specific $^{19}$F incorporation into uridine residues.** 1D $^{19}$F NMR spectra showing selective labeling efficiency at positions **(A)** U5, **(B)** U10, **(C)** U18, and **(D)** U25 within the RNA construct. Percentages indicate the relative population of each fluorinated species at the corresponding site.

A

| | H-REMD | roundtrips | % acceptance | time (μs) |
|---|---|---|---|---|
| apo | xOL3 | 83-2654 | 35-40 | 3 |
| | DESRES | 742-3814 | 33-39 | 3 |
| w/ C1 | A11-U33 | 2-223 | 14-20 | 1.5 |
| | A12-U33 | 2-1399 | 13-21 | 1.5 |
| | Mutant | 60-3482 | 29-34 | 1.5 |

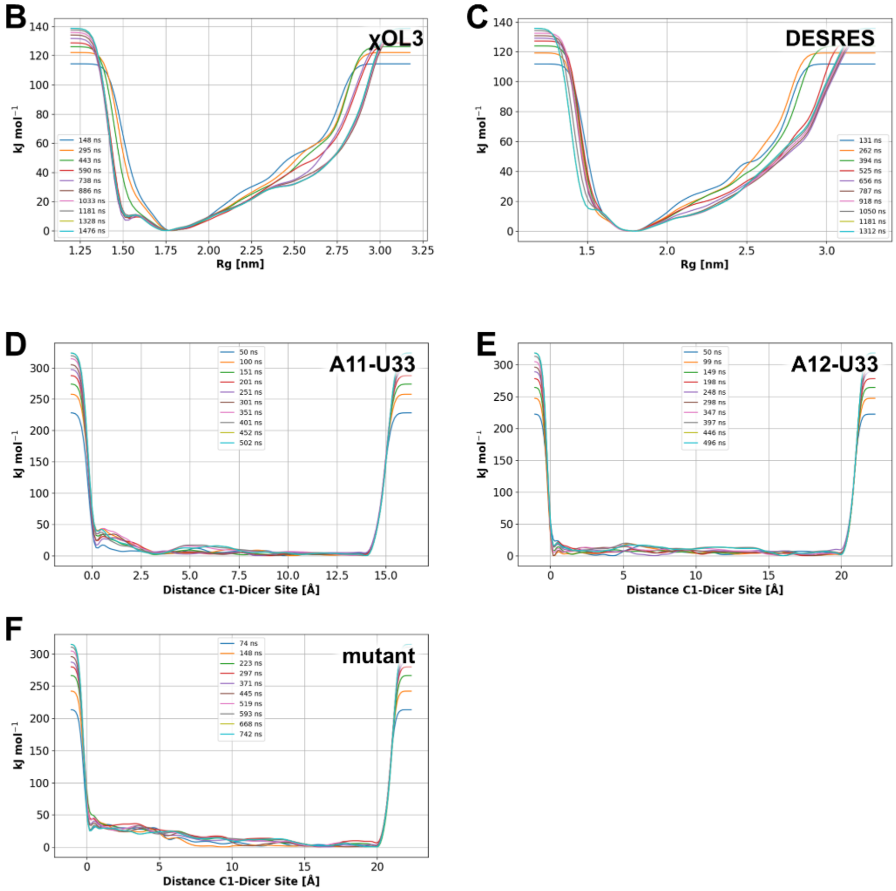

**Figure S8. Convergence assessment of enhanced-sampling simulations. (A)** Summary of H-REMD performance including roundtrips, exchange acceptance rates, and simulation times. **(B-C)** Convergence of metadynamics simulations for apo pre-miR-377 using radius of gyration as the collective variable. Overlapping block-wise free-energy profiles indicate converged sampling for both (B) xOL3 and (C) DESRES ensembles. **(D-F)** Block-wise free-energy profiles for holo systems (RNA-C1 complex) plotted versus distance between C1 and Dicer binding site for (D) A11-U33, (E) A12-U33, and (F) mutant constructs.

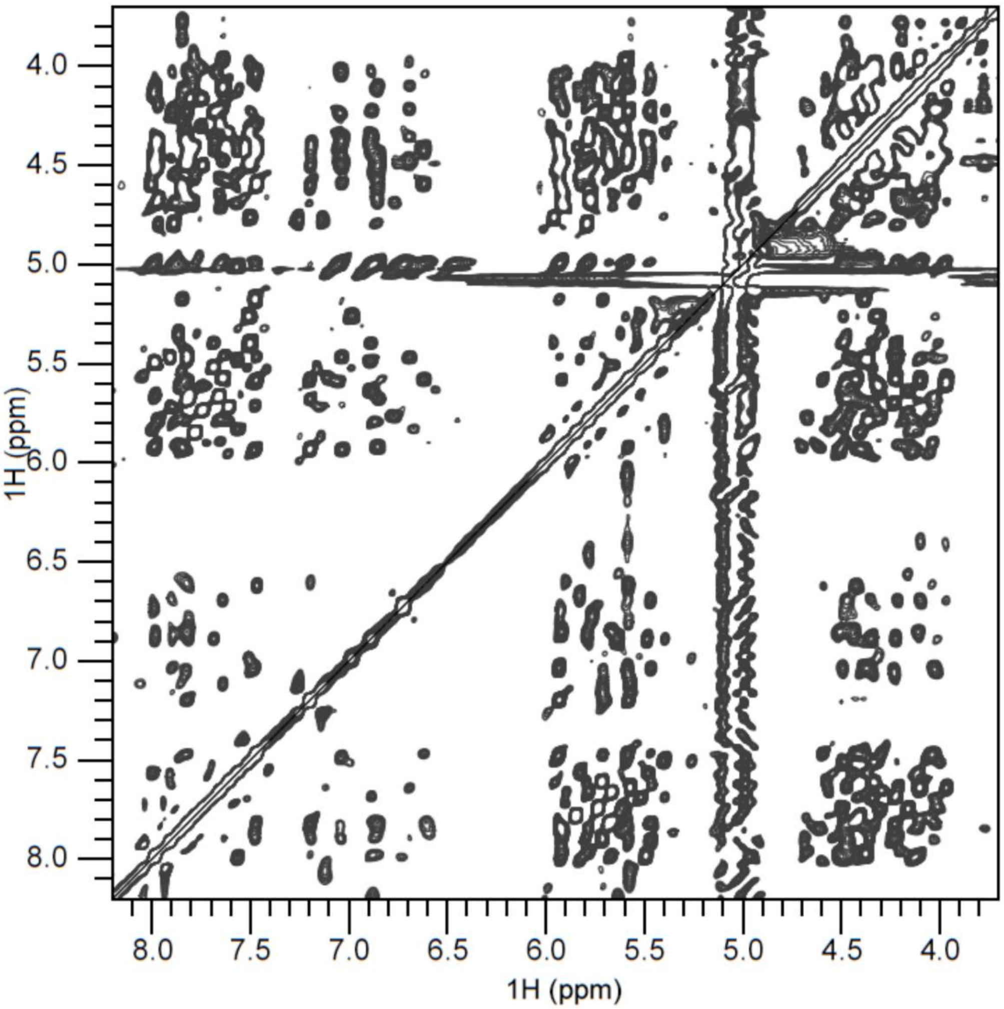

**Figure S9. 2D $^{1}H$-$^{1}H$ NOESY spectrum of pre-miR-377.** NOESY spectrum showing inter- and intra-nucleotide proton-proton correlations used to derive distance restraints for structure determination. Diagonal peaks correspond to identical protons, whereas off-diagonal cross peaks report short-range spatial contacts.

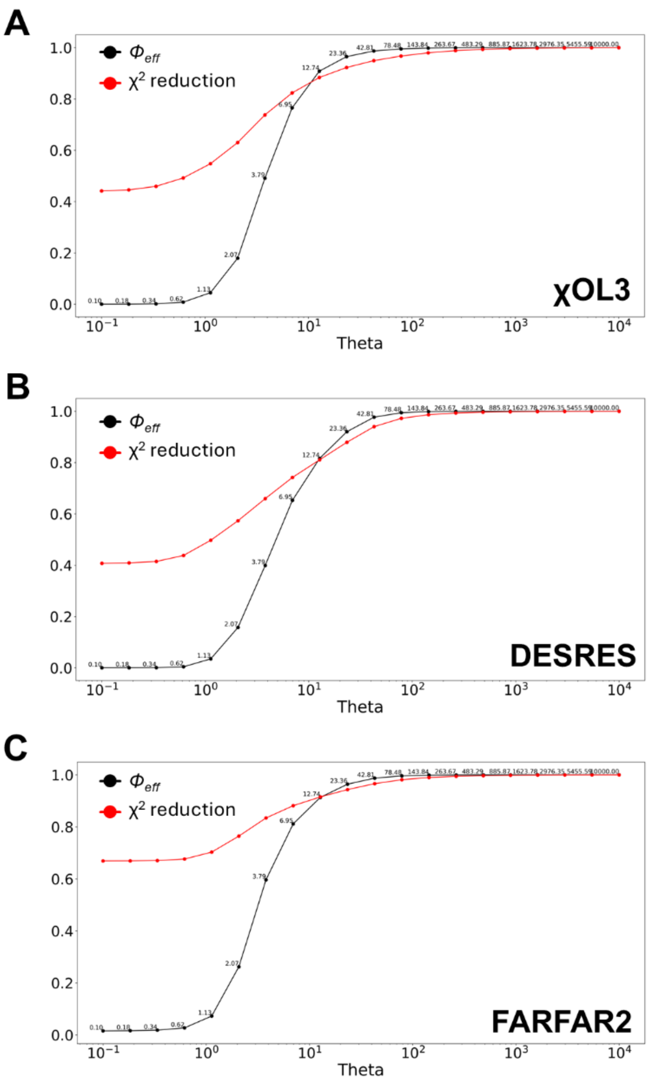

**Figure S10. Validation of structural ensembles of pre-miR-377 against NMR proton-proton distance restraints.** Reweighting analysis of *in silico* ensembles obtained from molecular dynamics simulations using **(A)** xOL3, **(B)** DESRES force fields, and **(C)** the FARFAR2 fragment assembly algorithm. For each model, proton-proton distances back-calculated from the simulations were reweighed across a range of $\theta$ (theta) values to improve agreement with experimental NMR restraints. The red curves indicate the $\chi^2$ reduction relative to the prior ensemble, while the black curves show $\Phi_{eff}$, representing the effective fraction of frames contributing after reweighting. To minimize model-specific overfitting and allow unbiased comparison among ensembles, a fixed value of $\theta = 1.13$ was applied across all modeling approaches.

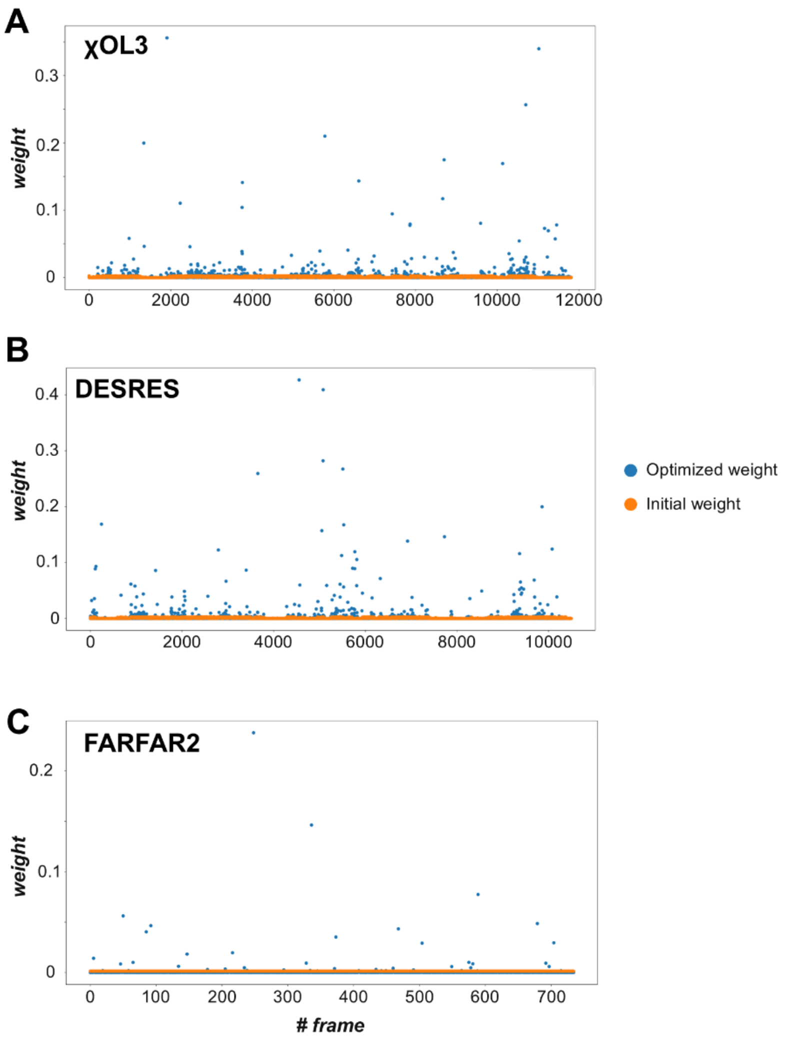

**Figure S11. Structural weights obtained from ensemble reweighting of pre-miR-377 models. (A-C)** Distribution of weights upon reweighting of ensembles generated using MD simulations with (A) xOL3, (B) DESRES force fields, and (C) FARFAR2 modeling approach. Each point represents the weight assigned to an individual structure after reweighting against experimental NMR proton-proton distance restraints. Orange dots indicate the prior distribution.

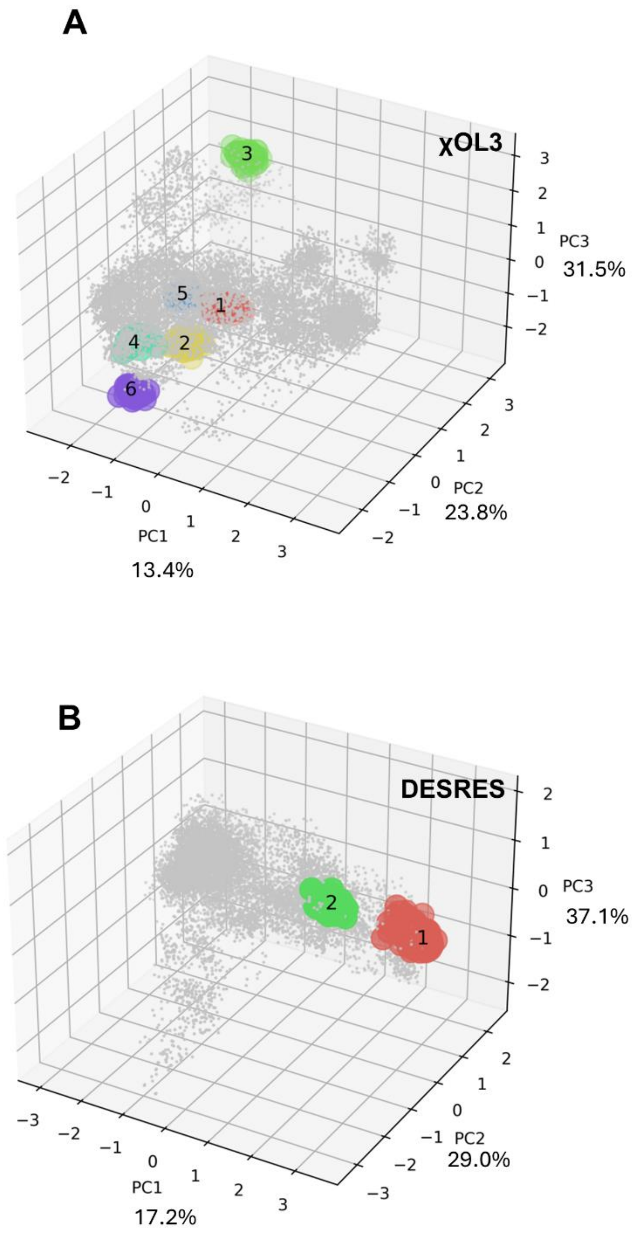

**Figure S12. Projection of structural clusters onto principal component space. (A-B)** Principal component analysis (PCA) of the MD-generated ensemble using (A) χOL3 and (B) DESRES force fields. Individual conformations are shown as gray points, and structures belonging to same cluster are colored accordingly.

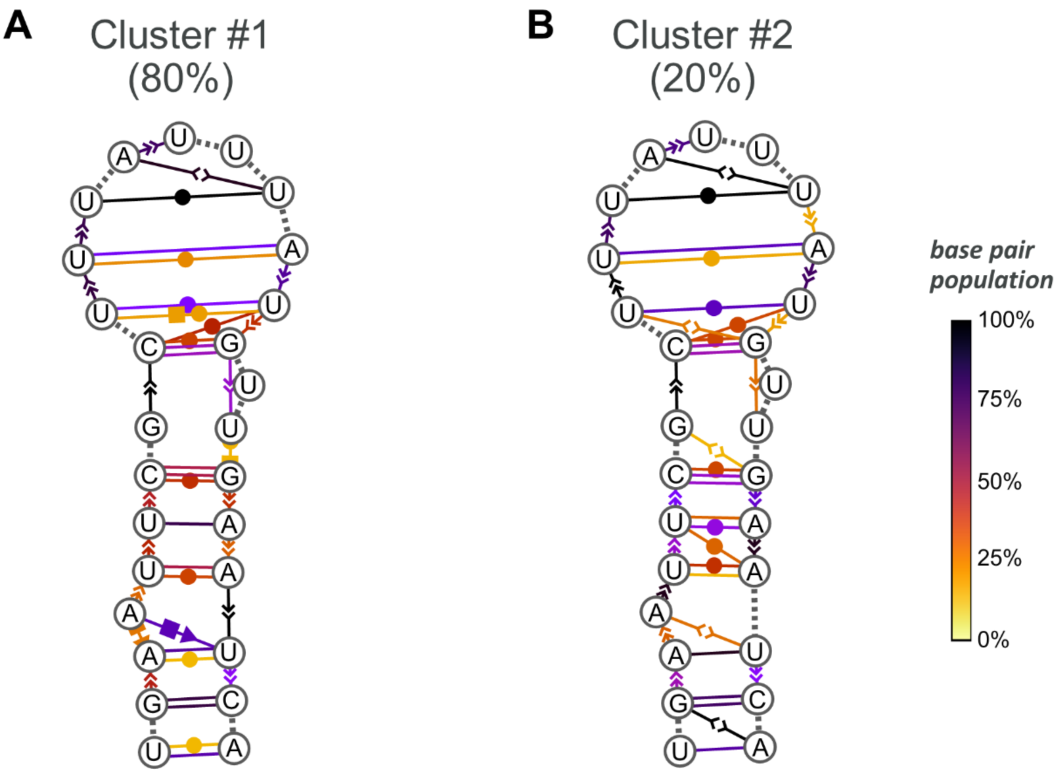

**Figure S13. Structural clustering of pre-miR-377 ensemble sampled with MD simulations using DESRES force field.** The secondary structure of each cluster is reported, and base-pair interactions are annotated using the Leontis-Westhof classification. The color gradient of connecting lines (yellow to black) represents the population density of each interaction, with darker purple indicating higher population. The percentage shown for each cluster indicates its relative population among all clustered structures.

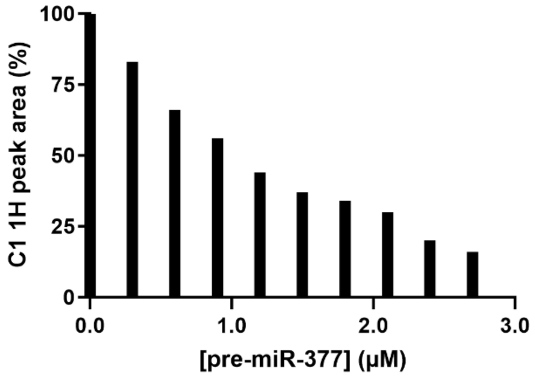

**Figure S14. CPMG titration of C1 reveals direct interaction with pre-miR-377.** $^1$H C1 peak areas decrease progressively with increasing pre-miR-377 concentration (0-3 μM), indicating binding-induced line broadening characteristic of intermediate exchange on the NMR timescale.

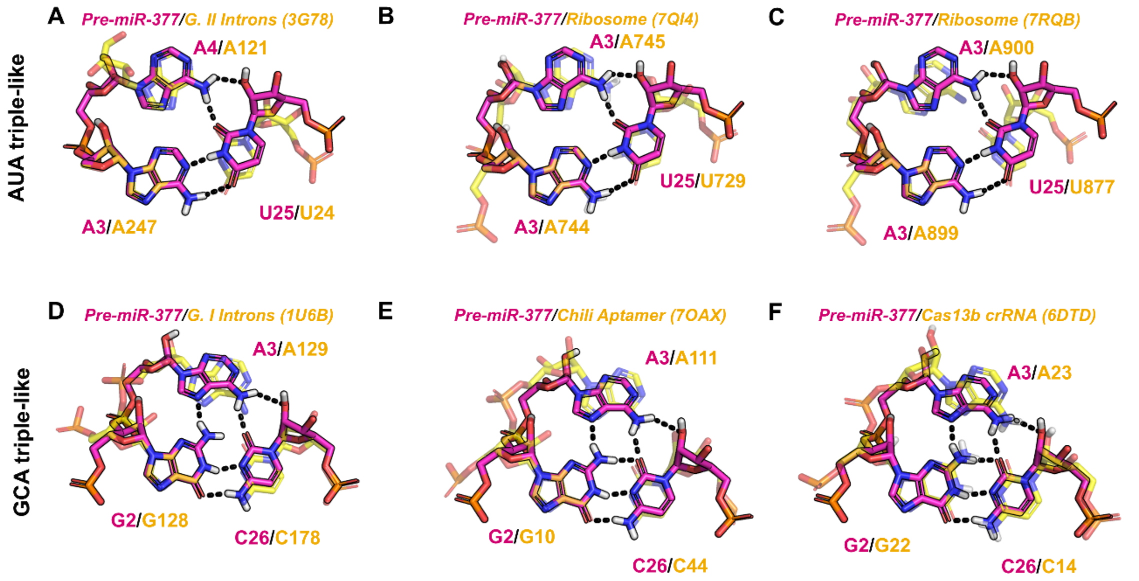

**Figure S15. Structural comparison of pre-miR-377 modeled base triples with experimentally determined AUA- and GCA-type triples from diverse regulatory RNAs. (A-C)** Superposition of the modeled AUA triple in pre-miR-377 (magenta) with corresponding AUA triples from Group II intron (PDB: 3G78, A247-U24-A121), mitochondrial ribosome (PDB: 7QI4, A744-U729-A745), and 70S ribosomal unit (PDB: 7RQB, A899-U877-A900; yellow). **(D-F)** Superposition of the modeled GCA triple in pre-miR-377 (magenta) with analogous GCA triples from Group I intron (PDB: 1U6B, G128-C178-A129), Chili RNA aptamer (PDB: 7OAX, G10-C44-A111), and Cas13b crRNA (PDB: 6DTD, G22-C14-A23; yellow). Hydrogen bonds are shown as black dashed lines. Pre-miR-377 bases are depicted as magenta sticks; reference RNA bases from other systems are shown as semi-transparent yellow sticks.

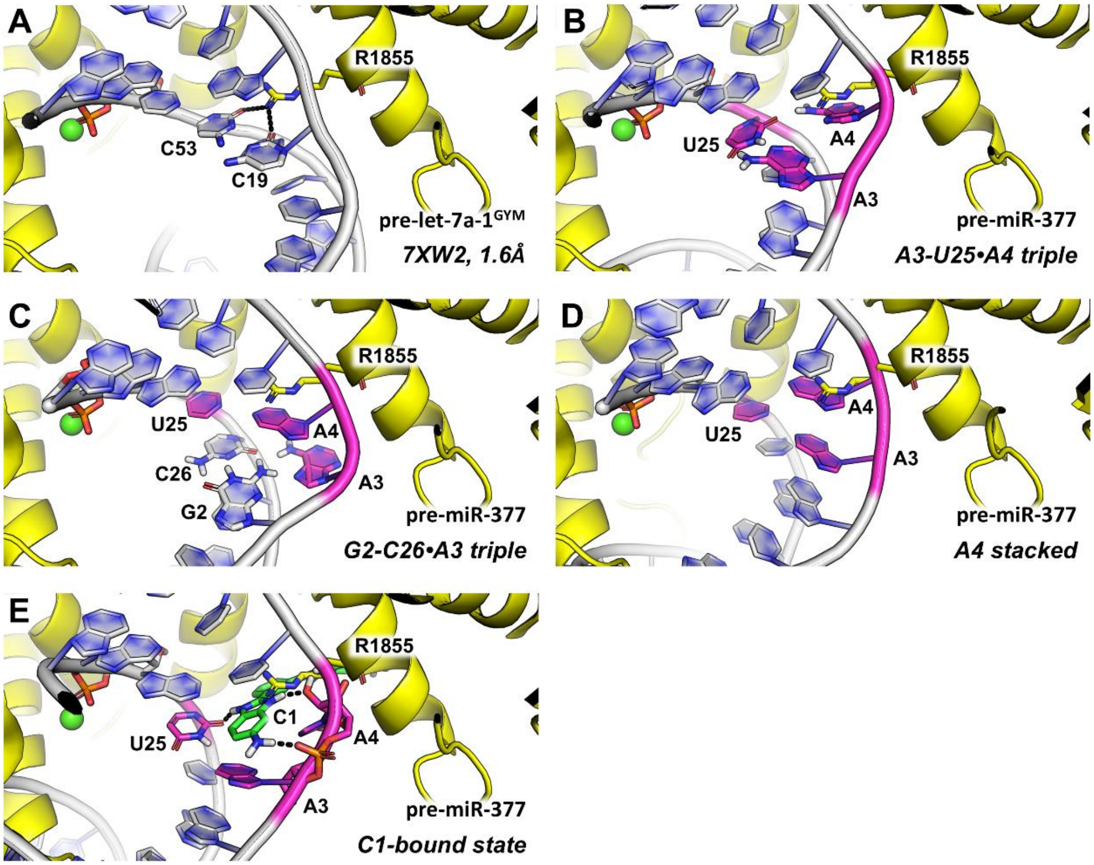

**Figure S16. Structural comparison of Dicer bound to diverse pre-miRNA substrates. (A)** Experimental structure of human Dicer bound to *pre-let-7a-1$^{GYM}$* (PDB ID: 7XW2, 1.6 Å), showing the positioning of residue R1855 (yellow) relative to the RNA duplex (gray). **(B-E)** Structural models of Dicer bound to pre-miR-377 conformations generated in this study. Regardless the RNA conformation, i.e., A3-U25•A4 triple (B), G2-C26•A3 triple (C), A4 stacked (D), or C1-bound state (E), the key residue R1855 clashes with the RNA substrate, suggesting that conformational reorganization of one or both binding partners is needed to guarantee RNA-protein recognition. In all panels, Dicer and RNA are depicted as cartoons (yellow and white, respectively). RNA bases involved in key interactions are shown as sticks, and nucleotides responsible for pre-miR-377 A-bulge shuffling are highlighted in magenta. C1 is shown as green sticks. The catalytic ion is represented as a green sphere, and the phosphate at the cleavage site is shown as sticks.

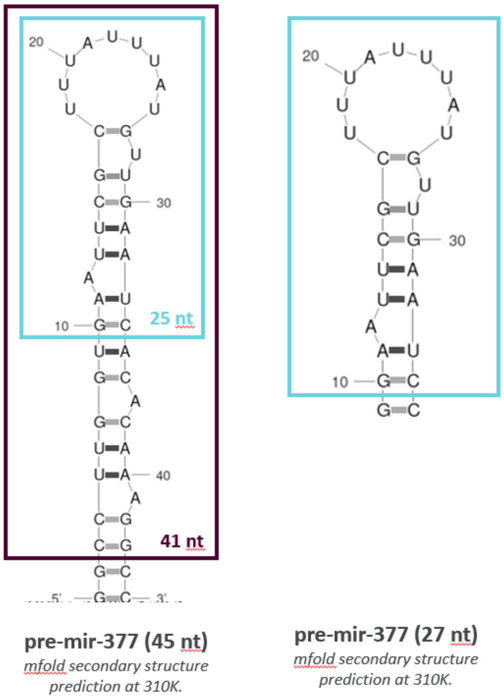

**Figure S17. Secondary structure of pre-miR-377 constructs.**

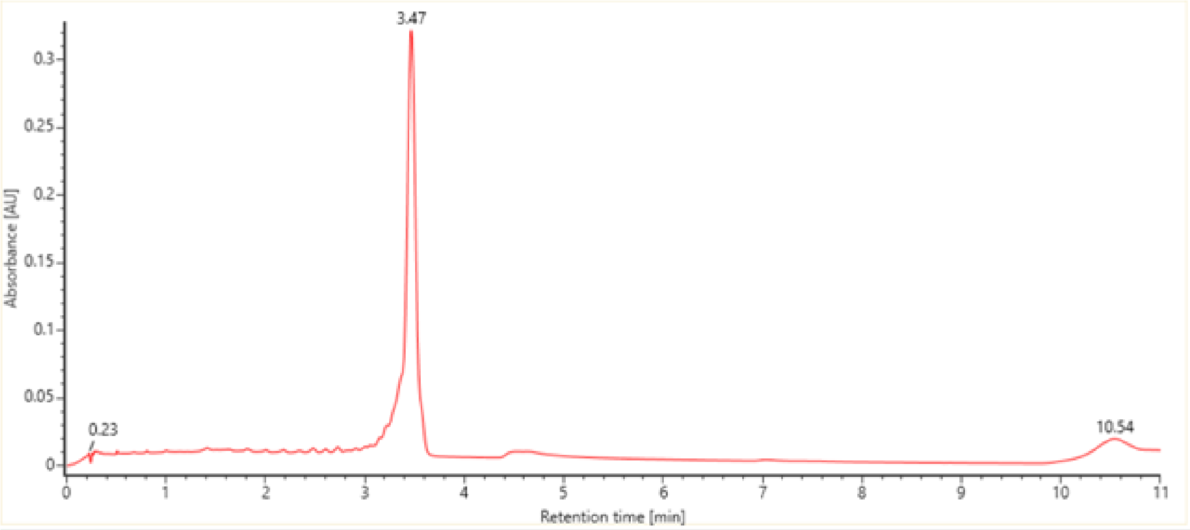

**Figure S18. Purified rFU5**. UGAA**U**UCGCUUUAUUUAUGUUGAAUCA. HRMS (ESI-) calculated 8536.9905, observed deconvoluted (BayerSpray) 8537.2767. UV Purity at 260 nm: 83%. yield 0.88%

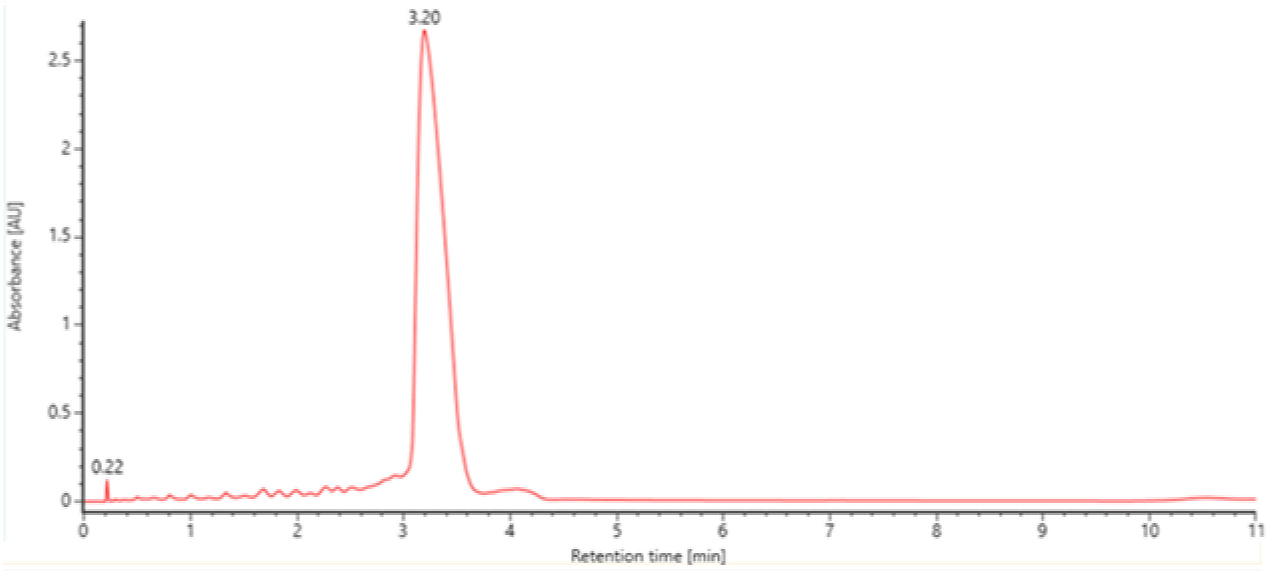

**Figure S19. Purified rFU10**. UGAAUUCGC**U**UUAUUUAUGUUGAAUCA. HRMS (ESI-) calculated 8536.9905, observed deconvoluted (BayerSpray) 8537.2041. UV Purity at 260 nm: 87%. yield 3.15%

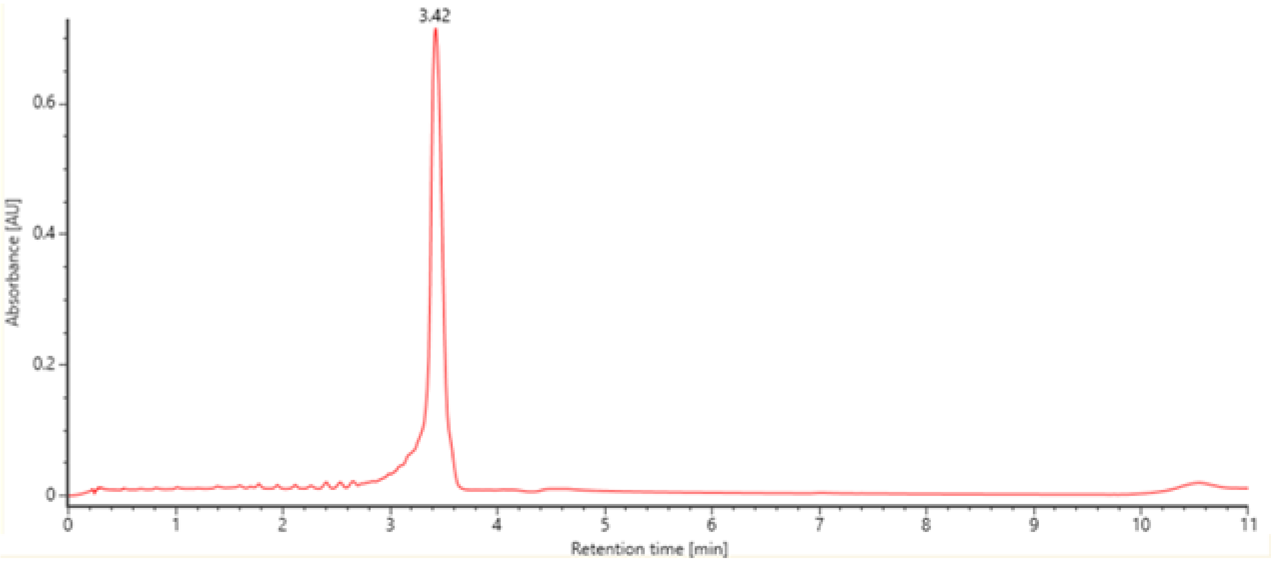

**Figure S20. Purified rFU18.** UGAAUUCGCUUUAUUUA**U**GUUGAAUCA. HRMS (ESI-) calculated 8536.9905, observed deconvoluted (BayerSpray) 8537.2692. UV Purity at 260 nm: 81%. yield 2.33%.

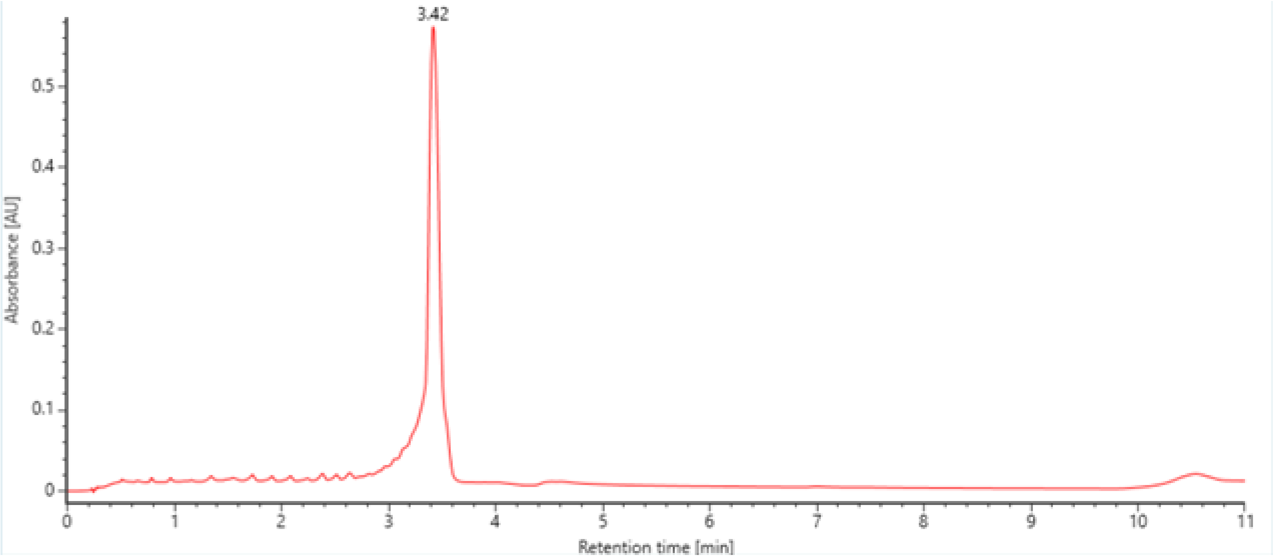

**Figure S21. Purified rFU25.** UGAAUUCGCUUUAUUUAUGUUGAA**U**CA. HRMS (ESI-) calculated 8536.9905, observed deconvoluted (BayerSpray) 8537.2678. UV Purity at 260 nm:81%. yield 0.78%

| pre-miR-377 construct | NMR isotope labels |
|---|---|
| **G**GAAUUCGC**U**UUAU**U**UAU**G**UUGAAUCC | |
| GGAAUUCGCU**U**UAUU**U**AU**G**UUGAAUCC | |
| G**G**AAUUCGCUUUAUUUAUGUUGAA**U**CC | |
| GGAAUUCGCUUUAUUUAU**G**UUGAAUCC | |
| GGAAU**U**CGCUUUAUUUAUGUU**G**AAUCC | $^{15}$N - H3N3, H1N1 |
| G**G**AAUUCGCUUUAUUUAUGUUGAA**U**CC | |
| GGAAU**U**CGCUUUAUUUAUGUU**G**AAUCC | |
| GGAAUUCGCU**U**UAUU**U**AUGUUGAAUCC | |
| GGAAUUCGC**U**UUAU**U**UAUGUUGAAUCC | |
| **G**GAAUUCGCUUU**A**U**U**UAUGUUGAAUCC | |
| GGAAUUCGCU**U**UAUUU**A**UGUU**G**AAUCC | |
| GGA**A**UU**C**GCUUUAUUUAU**G**UUGAAUCC | $^{13}$C - H8C8, H2C2, H6C6 |
| GGAAUUCGCUUUAUUUAUGUUG**A**A**U**CC | |
| GGAAU**U**CGCUUUAUUUAUGUU**G**A**A**UCC | |
| GGA**A**UUCGCUUUAUUUAUGCCUAAUCC | $^{19}$F - 2'OH |
| GGAAUUCGCUUUAUUUAUGCCUAA**U**CC | |
| UGAA**U**UCGCUUUAUUUAUGCCUAAUCA | |
| UGAAUUCGC**U**UUAUUUAUGCCUAAUCA | $^{19}$F - H5C5 |
| UGAAUUCGCUUUAUUUA**U**GCCUAAUCA | |
| UGAAUUCGCUUUAUUUAUGCCUAA**U**CA | |
| GGAAUUCGCUUUAUUUAUGCCUAAUCC | $^{15}$N and $^{13}$C fully labeled |

**Table S1. Sequences and isotope-labeling schemes for pre-miR-377 constructs used in this study.** All RNA constructs prepared for NMR experiments are listed, with the labeled nucleotides highlighted in bold within each sequence. The right column specifies the corresponding isotope-labeling scheme, including site-specific $^{15}$N, $^{13}$C, and $^{19}$F labeling at designated positions, as well as the fully $^{15}$N- and $^{13}$C-labeled sample.

| pos. #1 | nt. #1 | H #1 | pos. #2 | nt. #2 | H #2 | H-H distance | error |
|---|---|---|---|---|---|---|---|
| 2 | G | H1 | 26 | C | H2' | 4.57 | 0.3 |
| 2 | G | H1 | 26 | C | H5 | 5.16 | 0.39 |
| 2 | G | H1 | 26 | C | H6 | 5.44 | 1.9 |
| 4 | A | H2 | 24 | A | H61 | 4.68 | 0.93 |
| 4 | A | H2 | 4 | A | H1' | 4.39 | 0.54 |
| 4 | A | H2 | 4 | A | H61 | 4.63 | 0.53 |
| 4 | A | H2 | 5 | U | H5 | 4.54 | 0.53 |
| 4 | A | H8 | 4 | A | H1' | 3.41 | 0.47 |
| 4 | A | H8 | 4 | A | H3' | 3.06 | 0.54 |
| 4 | A | H8 | 4 | A | H5' | 3.49 | 0.52 |
| 4 | A | H8 | 4 | A | H61 | 4.64 | 1.21 |
| 4 | A | H8 | 5 | U | H5 | 3.85 | 0.5 |
| 4 | A | H8 | 5 | U | H6 | 4.19 | 0.8 |
| 5 | U | H1' | 5 | U | H3' | 3.85 | 0.4 |
| 5 | U | H5 | 6 | U | H5 | 3.7 | 0.59 |
| 5 | U | H6 | 4 | A | H3' | 4.11 | 0.5 |
| 5 | U | H6 | 5 | U | H2' | 3.4 | 0.53 |
| 5 | U | H6 | 5 | U | H5 | 2.99 | 0.47 |
| 5 | U | H6 | 5 | U | H5' | 3.76 | 0.6 |
| 6 | U | H1' | 6 | U | H2' | 2.96 | 0.92 |
| 6 | U | H1' | 6 | U | H6 | 3.22 | 0.56 |
| 6 | U | H3 | 22 | G | H1 | 3.97 | 0.3 |
| 6 | U | H3 | 23 | A | H2 | 3.21 | 1 |
| 6 | U | H3 | 7 | C | H41 | 4.93 | 1.8 |
| 6 | U | H3 | 7 | C | H5 | 4.68 | 1.68 |
| 6 | U | H5 | 5 | U | H3' | 3.13 | 0.55 |
| 6 | U | H5 | 5 | U | H5 | 3.8 | 0.6 |
| 6 | U | H6 | 6 | U | H2' | 3.47 | 0.57 |
| 6 | U | H6 | 6 | U | H3' | 2.86 | 0.48 |
| 6 | U | H6 | 6 | U | H5' | 3.46 | 0.47 |
| 7 | C | H41 | 7 | C | H5 | 3.36 | 0.48 |
| 7 | C | H42 | 23 | A | H2 | 5.5 | 1.01 |
| 7 | C | H42 | 7 | C | H5 | 4.34 | 1.82 |
| 7 | C | H5 | 6 | U | H6 | 3.91 | 0.6 |
| 7 | C | H6 | 7 | C | H41 | 3.93 | 0.41 |
| 7 | C | H6 | 7 | C | H5 | 2.89 | 0.58 |
| 8 | G | H1 | 22 | G | H8 | 4.49 | 0.3 |
| 8 | G | H1 | 9 | C | H41 | 4.91 | 0.3 |
| 8 | G | H1 | 9 | C | H42 | 4.69 | 0.3 |
| 8 | G | H8 | 8 | G | H1' | 3.13 | 0.45 |
| 8 | G | H8 | 8 | G | H5' | 3.54 | 0.52 |
| 8 | G | H8 | 9 | C | H5 | 4.16 | 0.84 |
| 9 | C | H1' | 9 | C | H2' | 2.92 | 0.45 |
| 9 | C | H41 | 10 | U | H5 | 4.7 | 0.43 |
| 9 | C | H41 | 9 | C | H5 | 3.35 | 0.41 |
| 9 | C | H42 | 9 | C | H5 | 3.85 | 1.23 |
| 9 | C | H5 | 9 | C | H2' | 4.16 | 1.76 |
| 9 | C | H5 | 9 | C | H3' | 4.08 | 1.73 |
| 9 | C | H6 | 10 | U | H5 | 4.05 | 0.41 |
| 9 | C | H6 | 8 | G | H1' | 4.37 | 0.52 |
| 9 | C | H6 | 9 | C | H3' | 3.24 | 0.47 |
| 9 | C | H6 | 9 | C | H41 | 4.29 | 0.38 |
| 9 | C | H6 | 9 | C | H5 | 3.04 | 0.44 |
| 10 | U | H6 | 10 | U | H2' | 3.35 | 0.47 |
| 10 | U | H6 | 10 | U | H3' | 2.97 | 0.43 |
| 10 | U | H6 | 10 | U | H5 | 2.94 | 0.47 |
| 10 | U | H6 | 10 | U | H5' | 3.89 | 0.52 |
| 10 | U | H6 | 9 | C | H41 | 5.8 | 0.43 |
| 10 | U | H6 | 9 | C | H5 | 5.55 | 1.75 |
| 10 | U | H5 | 9 | C | H5 | 3.85 | 0.72 |
| 11 | U | H6 | 11 | U | H5 | 2.95 | 0.74 |
| 12 | U | H6 | 12 | U | H3' | 2.6 | 0.43 |
| 12 | U | H6 | 12 | U | H5 | 2.45 | 0.38 |
| 13 | A | H1' | 13 | A | H2' | 2.94 | 0.4 |
| 13 | A | H1' | 13 | A | H3' | 2.95 | 0.88 |
| 13 | A | H1' | 16 | U | H5 | 3.21 | 0.56 |
| 13 | A | H1' | 16 | U | H5 | 3.21 | 0.56 |
| 13 | A | H1' | 17 | A | H5" | 3.94 | 0.49 |
| 13 | A | H2 | 13 | A | H62 | 4.02 | 0.76 |
| 13 | A | H2 | 17 | A | H5" | 3.49 | 0.58 |
| 13 | A | H62 | 12 | U | H1' | 5.1 | 0.43 |
| 13 | A | H62 | 14 | U | H5 | 5.01 | 0.43 |
| 13 | A | H8 | 13 | A | H1' | 3 | 0.45 |
| 13 | A | H8 | 13 | A | H2' | 3.12 | 0.54 |
| 13 | A | H8 | 13 | A | H3' | 2.61 | 0.44 |
| 13 | A | H8 | 13 | A | H61 | 5.22 | 0.42 |
| 13 | A | H8 | 13 | A | H62 | 4.86 | 0.7 |
| 13 | A | H8 | 14 | U | H6 | 4.32 | 0.85 |
| 13 | A | H8 | 16 | U | H6 | 4.33 | 0.78 |
| 14 | U | H1' | 13 | A | H2' | 3.06 | 0.45 |
| 14 | U | H1' | 14 | U | H2' | 2.88 | 0.55 |
| 14 | U | H1' | 14 | U | H4' | 2.97 | 0.47 |
| 14 | U | H6 | 13 | A | H1' | 4.56 | 0.55 |
| 14 | U | H6 | 13 | A | H61 | 5.57 | 0.45 |
| 14 | U | H6 | 14 | U | H1' | 3.46 | 0.54 |
| 14 | U | H6 | 14 | U | H2' | 3.04 | 0.46 |
| 14 | U | H6 | 14 | U | H3' | 2.83 | 0.39 |
| 14 | U | H6 | 14 | U | H5 | 2.65 | 0.44 |
| 14 | U | H6 | 14 | U | H5' | 3.71 | 0.6 |
| 15 | U | H1' | 15 | U | H2' | 2.87 | 0.96 |
| 15 | U | H1' | 15 | U | H3' | 3.2 | 0.59 |
| 15 | U | H1' | 15 | U | H4' | 3.27 | 0.7 |
| 15 | U | H2' | 15 | U | H3' | 2.18 | 0.47 |
| 15 | U | H3' | 15 | U | H4' | 2.35 | 0.49 |
| 15 | U | H6 | 15 | U | H1' | 3.44 | 0.57 |
| 15 | U | H6 | 15 | U | H3' | 3.09 | 0.49 |
| 15 | U | H6 | 15 | U | H5 | 2.62 | 0.44 |
| 15 | U | H6 | 15 | U | H5' | 3.74 | 0.62 |
| 15 | U | H6 | 17 | A | H1' | 4.61 | 0.58 |
| 16 | U | H1' | 12 | U | H6 | 3.2 | 0.56 |
| 16 | U | H1' | 16 | U | H2' | 3.08 | 0.55 |
| 16 | U | H6 | 16 | U | H1' | 3.15 | 0.39 |
| 16 | U | H6 | 16 | U | H3' | 2.7 | 0.42 |
| 16 | U | H6 | 16 | U | H5 | 2.57 | 0.4 |
| 17 | A | H1' | 17 | A | H2' | 2.98 | 0.86 |
| 17 | A | H1' | 17 | A | H3' | 3.12 | 1.06 |
| 17 | A | H1' | 17 | A | H4' | 3.25 | 0.72 |
| 17 | A | H2 | 12 | U | H1' | 3.09 | 0.44 |
| 17 | A | H2 | 13 | A | H1' | 3.76 | 0.47 |
| 17 | A | H2 | 14 | U | H1' | 2.99 | 0.42 |
| 17 | A | H2 | 16 | U | H1' | 3.85 | 0.63 |
| 17 | A | H2 | 17 | A | H1' | 3.89 | 0.9 |
| 17 | A | H8 | 12 | U | H1' | 3.89 | 0.3 |
| 17 | A | H8 | 12 | U | H5 | 3.94 | 0.66 |
| 17 | A | H8 | 12 | U | H6 | 4.1 | 0.47 |
| 17 | A | H8 | 15 | U | H6 | 4.75 | 0.38 |
| 17 | A | H8 | 16 | U | H1' | 3.8 | 0.56 |
| 17 | A | H8 | 17 | A | H1' | 3.1 | 0.49 |
| 17 | A | H8 | 17 | A | H2' | 3 | 0.43 |
| 17 | A | H8 | 17 | A | H3' | 2.94 | 0.53 |
| 17 | A | H8 | 18 | U | H6 | 4.56 | 1.01 |
| 18 | U | H1' | 17 | A | H2 | 3.38 | 0.57 |
| 18 | U | H1' | 18 | U | H2' | 3.39 | 0.75 |
| 18 | U | H1' | 18 | U | H3' | 3.85 | 0.91 |
| 18 | U | H1' | 18 | U | H4' | 3.35 | 0.69 |
| 18 | U | H5 | 17 | A | H2' | 3.56 | 0.58 |
| 18 | U | H5 | 17 | A | H8 | 4.11 | 0.62 |
| 18 | U | H6 | 17 | A | H1' | 4.07 | 0.69 |
| 18 | U | H6 | 18 | U | H3' | 2.78 | 0.42 |
| 18 | U | H6 | 18 | U | H4' | 4.12 | 0.41 |
| 18 | U | H6 | 18 | U | H5' | 3.76 | 0.72 |
| 18 | U | H6 | 18 | U | H5" | 3.75 | 1.02 |
| 19 | G | H1 | 10 | U | H5 | 5.37 | 1.15 |
| 19 | G | H1 | 10 | U | H6 | 5.37 | 1.48 |
| 19 | G | H1 | 9 | C | H42 | 3.72 | 1.1 |
| 19 | G | H1 | 9 | C | H5 | 4.82 | 1.47 |
| 19 | G | H1 | 9 | C | H6 | 5.43 | 1.45 |
| 19 | G | H8 | 19 | G | H5' | 3.78 | 0.6 |
| 19 | G | H8 | 19 | G | H5" | 3.7 | 0.59 |
| 22 | G | H1 | 22 | G | H1' | 3.82 | 1.76 |
| 22 | G | H1 | 23 | A | H2 | 4.21 | 1.57 |
| 22 | G | H1 | 7 | C | H41 | 3.87 | 1.83 |
| 22 | G | H1 | 7 | C | H42 | 3.63 | 1.64 |
| 22 | G | H1 | 8 | G | H1 | 4.99 | 0.3 |
| 22 | G | H1 | 8 | G | H21 | 5.24 | 2.26 |
| 22 | G | H1 | 8 | G | H8 | 4.74 | 1.93 |
| 22 | G | H1' | 22 | G | H2' | 2.64 | 0.39 |
| 22 | G | H1' | 23 | A | H1' | 4.76 | 0.53 |
| 22 | G | H21 | 22 | G | H1' | 3.54 | 0.39 |
| 22 | G | H21 | 23 | A | H1' | 3.61 | 0.39 |
| 22 | G | H8 | 22 | G | H1' | 3.77 | 0.68 |
| 23 | A | H2 | 22 | G | H21 | 5.94 | 2.36 |
| 23 | A | H2 | 23 | A | H1' | 4.15 | 0.5 |
| 23 | A | H2 | 24 | A | H1' | 3.07 | 0.62 |
| 23 | A | H2 | 7 | C | H1' | 3.26 | 0.56 |
| 24 | A | H1' | 24 | A | H2' | 2.58 | 0.39 |
| 24 | A | H1' | 25 | U | H1' | 4.33 | 0.52 |
| 24 | A | H8 | 24 | A | H5' | 3.75 | 0.49 |
| 24 | A | H8 | 24 | A | H5" | 4.02 | 0.59 |
| 24 | A | H8 | 25 | U | H5 | 3.98 | 0.52 |
| 25 | U | H1' | 25 | U | H3' | 3.28 | 1.31 |
| 25 | U | H1' | 25 | U | H4' | 3.15 | 0.68 |
| 25 | U | H1' | 25 | U | H5 | 4.84 | 0.55 |
| 25 | U | H3 | 4 | A | H2 | 4.68 | 0.48 |
| 25 | U | H6 | 25 | U | H1' | 3.78 | 0.59 |
| 25 | U | H6 | 25 | U | H2' | 3.23 | 0.46 |
| 25 | U | H6 | 25 | U | H4' | 4.17 | 0.51 |
| 25 | U | H6 | 25 | U | H5 | 3.02 | 0.52 |
| 26 | C | H1' | 26 | C | H2' | 2.66 | 0.51 |
| 26 | C | H1' | 26 | C | H3' | 3.14 | 0.62 |
| 26 | C | H42 | 26 | C | H5 | 3.55 | 0.37 |
| 26 | C | H6 | 25 | U | H1' | 3.92 | 0.78 |
| 26 | C | H6 | 26 | C | H2' | 3.2 | 0.49 |
| 26 | C | H6 | 26 | C | H3' | 2.93 | 0.44 |
| 26 | C | H6 | 26 | C | H41 | 4.51 | 0.91 |
| 26 | C | H6 | 26 | C | H42 | 4.2 | 0.57 |
| 26 | C | H6 | 26 | C | H5 | 2.91 | 0.55 |
| 26 | C | H6 | 26 | C | H5' | 3.35 | 0.41 |

**Table S2. Experimental NMR-derived proton–proton distance restraints used for structural validation.** Each row lists an observed H-H cross-peak assigned between two nuclei in pre-miR-377. Columns report: the nucleotide positions involved (pos. #1 and pos. #2), the corresponding nucleotide identities (nt. #1 and nt. #2), the specific protons assigned (H #1 and H #2), the extracted interproton distance (Å), and the associated experimental uncertainty.

| RNA type | Res. (Å) | PDBid | Nt #1 | Nt #2 | Nt #3 | Sequence order | LW 1- 2 | LW 1-3 | LW 2-3 |
|---|---|---|---|---|---|---|---|---|---|
| **AUA triples** | | | | | | | | | |
| mitochondrial ribosome | 2.2 | 7QI4 | A744 | U729 | A745 | A←U→A | cWW | | tSH |
| 70S ribosomal unit | 2.5 | 7RQB | A899 | U877 | A900 | A←U→A | cWW | | tSH |
| Group II Intron | 2.9 | 4FAR | A247 | U24 | A121 | A←U→A | cWW | | tSH |
| Hibernating ribosome | 2.9 | 8FMW | A979 | U958 | A980 | A←U→A | cWW | | tSH |
| 55S mitochondrial ribosome | 3.0 | 6YDP | A1208 | U1194 | A1209 | A←U→A | cWW | | tSH |
| Group II Intron | 3.0 | 8T2S | A255 | U24 | A124 | A←U→A | cWW | ncSH | tSH |
| **GCA triples** | | | | | | | | | |
| Cas13b CRISPR RNA (crRNA) | 1.7 | 6DTD | G22 | C16 | A23 | G←C→A | cWW | ncSH | tSH |
| mitochondrial ribosome | 2.2 | 7QI4 | G1355 | C1209 | A1356 | G←C→A | cWW | cSH | tSH |
| Chili RNA aptamer | 2.2 | 7OAX | G10 | C44 | A11 | G→C←A | cWW | cSH | tSH |
| ydaO riboswitch | 2.7 | 4QLM | G40 | C91 | A95 | G→C→A | cWW | ncSH | tSH |
| 70S ribosomal unit | 2.8 | 7JIL | G1274 | C1260 | A1275 | G←C→A | cWW | | tSH |
| chloroplast ribosome | 3.0 | 6ERI | G1611 | C1442 | A1612 | G←C→A | cWW | cSH | tSH |
| group I intron | 3.1 | 1U6B | G128 | C178 | A129 | G←C→A | cWW | cSH | tSH |

**Table S3. Summary of RNA triple base interactions observed in available high-resolution structures.** Examples of AUA and GCA base triples identified from RNA-containing structures in the Protein Data Bank (PDB). The table lists the RNA type, structural resolution, PDB ID, and the nucleotide identities and numbering (Nt #1-3) involved in each triple. The sequence order column indicates the 5′→3′ base arrangement. LW 1-2, LW 1-3, and LW 2-3 denote the base-pairing geometries between the three nucleotides using the Leontis-Westhof (LW) nomenclature, where cWW = *cis* Watson-Crick/Watson-Crick, tSH = *trans* Sugar-Hoogsteen, cSH = *cis* Sugar-Hoogsteen, and ncSH = non-canonical Sugar-Hoogsteen.

| Spectro, 600 if not spec | spectrum | sample | temp | pp | td | ns | d1 | sw 1h | sw hetero atom | sw 1h 3D | o1 | o2 | o3 |
|---|---|---|---|---|---|---|---|---|---|---|---|---|---|
| | noesy D2O 50ms | unlabelled | 283 | noesyesgpph | 2048 / 512 | 128 | 1.5 | 5405 hz 9 ppm | 5402 hz 9ppm | | 2819hz 4.697ppm | | |
| | noesy D2O 150ms | unlabelled | 283 | noesyesgpph | 2048 / 448 | 128 | 1.5 | 5405 hz 9 ppm | 5402 hz 9ppm | | 2819hz 4.697ppm | | |
| | noesy H2O 100ms | unlabelled | 283 | noesyesfbgpph | 4096 / 736 | 256 | 1 | 13158 hz 22ppm | 5402 hz 9ppm | | 2815hz 4.69ppm | | |
| 800 | noesy H2O 150ms | unlabelled | 283 | na_noesy-jrecho-dec | 4096 / 384 | 384 | 1.5 | 16741 hz 21ppm | 16797 hz 21 ppm | | 3751 hz 4.69 ppm | | |
| | HC | A4 C7 G19 | 283 | sf_mehmqcgpph | 2048 / 256 | 128 | 0.3 | 8418 hz 14ppm | 5283hz 35ppm | | 2817hz 4.693ppm | 21432hz 142ppm | 9427hz 155ppm |
| | TOCSY 30ms | A4 C7 G19 | 283 | dipsi2esfbgpph | 2048 / 640 | 80 | 1 | 5411hz 9ppm | 3901hz 6.5ppm | | 2819hz 4.697ppm | 15244hz 101ppm | 5231hz 86ppm |
| | TOCSY 30ms | G2 A23 U25 | 283 | dipsi2esfbgpph | 2048 / 640 | 80 | 1 | 5411hz 9ppm | 3901hz 6.5ppm | | 2818hz 4.694ppm | 15244hz 101ppm | 5231hz 86ppm |
| | HC | G2 A23 U25 | 283 | sf_mehmqcgpph | 4096 / 512 | 128 | 0.3 | 8418 hz 14ppm | 5283hz 35ppm | | 2818hz 4.694ppm | 21432hz 142ppm | 9427hz 155ppm |
| | TOCSY 30ms | U6 G22 A24 | 283 | dipsi2esfbgpph | 2048 / 640 | 80 | 1 | 5411hz 9ppm | 3901hz 6.5ppm | | 2816hz 4.691ppm | 15244hz 101ppm | 5231hz 86ppm |
| | HC | U6 G22 A24 | 283 | sf_mehmqcgpph | 2048 / 256 | 128 | 0.3 | 8418 hz 14ppm | 5283hz 35ppm | | 2816hz 4.691ppm | 21432hz 142ppm | 9427hz 155ppm |
| | HN | U11 A17 G22 + U16 | 283 | sfhmqcf3gpph | 4096 / 384 | 128 | 0.25 | 13228hz 22ppm | 3650hz 60ppm | | 2816hz 4.691ppm | 22941hz 152ppm | 9731hz 160ppm |
| | HC | U11 A17 G22 + U16 | 283 | sf_mehmqcgpph | 4096 / 384 | 128 | 0.3 | 8418 hz 14ppm | 5283hz 35ppm | | 2816hz 4.691ppm | 21432hz 142ppm | 9427hz 155ppm |
| | TOCSY 30ms | U11 A17 G22 + U16 | 283 | dipsi2esfbgpph | 4096 / 640 | 128 | 1 | 5411hz 9ppm | 3901hz 6.5ppm | | 2816hz 4.691ppm | 15244hz 101ppm | 5231hz 86ppm |
| | HN | A13 U15 + G1 U10 | 283 | sfhmqcf3gpph | 4096 / 384 | 256 | 0.25 | 13228hz 22ppm | 3650hz 60ppm | | 2817hz 4.693ppm | 22941hz 152ppm | 9731hz 160ppm |
| | HC | A13 U15 + G1 U10 | 283 | sf_mehmqcgpph | 4096 / 384 | 128 | 0.3 | 8418 hz 14ppm | 5283hz 35ppm | | 2817hz 4.693ppm | 21432hz 142ppm | 9427hz 155ppm |

| | | | | | | | | | | | | | | |
|---|---|---|---|---|---|---|---|---|---|---|---|---|---|---|
| | TOCSY 30ms | A13 U15 + G1 U10 | 283 | dipsi2esfbgpph | 4096 / 512 | 128 | 1 | 5411hz 9ppm | 3901hz 6.5ppm | | 2817hz 4.693ppm | 15244hz 101ppm | 5231hz 86ppm | |
| | HN | U6 G22 A24 | 283 | sfhmqcf3gpph | 2048 / 128 | 128 | 0.25 | 13158hz 22ppm | 3649hz 60ppm | | 2821hz 4.7ppm | 22937hz 152ppm | 9730hz 160ppm | |
| | HN | G2 A23 U25 | 283 | sfhmqcf3gpph | 2048 / 128 | 128 | 0.25 | 13158hz 22ppm | 3649hz 60ppm | | 2821hz 4.7ppm | 22937hz 152ppm | 9730hz 160ppm | |
| | HN | A4 C7 G19 | 283 | sfhmqcf3gpph | 2048 / 128 | 128 | 0.25 | 13158hz 22ppm | 3649hz 60ppm | | 2821hz 4.7ppm | 22937hz 152ppm | 9730hz 160ppm | |
| | HC | unlabelled | 283 | sf_mehmqcgpph | 2048 / 512 | 768 | 0.25 | 8197hz 13.66ppm | 4528hz 30ppm | | 2816hz 4.691ppm | 21734hz 144ppm | 9427hz 155ppm | C862 |
| 800 | HC | 15N13C | 283 | hmqcetgpsi | 2048 / 320 | 16 | 1.5 | 8803hz 11ppm | 24138hz 120ppm | | 3759hz 4.7ppm | 20113hz 100ppm | 12158hz 150ppm | all C |
| | HC | 15N13C | 298 | hsqcetgpsisphd | 4096 / 512 | 64 | 1 | 5263hz 8.77ppm | 2264hz 15ppm | | 2816hz 4.693 | 13731hz 91ppm | 9729hz 160ppm | C1' |
| | HC | unlabelled | 298 | hsqcetgpsp | 4096 / 512 | 64 | 1 | 5882hz 9.8ppm | 3622hz 24ppm | | 2816hz 4.693ppm | 21427hz 142ppm | | C862 |
| | HN | 15N13C | 283 | sfhmqcf3gpph | 2048 / 512 | 320 | 0.25 | 14423hz 24ppm | 1521hz 25ppm | | 2821hz 4.7ppm | 21432hz 142ppm | 9427hz 155ppm | multiple UA |
| | HN | 15N13C | 283 | na_b_trhnncosygpph | 2048 / 512 | 448 | 0.25 | 13228hz 22ppm | 12166hz 200ppm | | 2821hz 4.7ppm | 22941hz 152ppm | 9731hz 160ppm | HNN COSY, multiple UAs |
| | noesy D2O 100ms | 15C13C | 283 | noesyhsqcgp3d | 2048 / 40 / 480 | 16 | 1.5 | 5882hz 9.8ppm | 6038hz 40ppm | 6002hz 10ppm | 2821hz 4.7ppm | 15847hz 105ppm | 6994hz 115ppm | 3D C5 |

**Table S4. Detailed NMR experimental parameters**: Spectral widths, time domain points, and acquisition parameters for each experiment type

**Phosphoramidite synthesis**

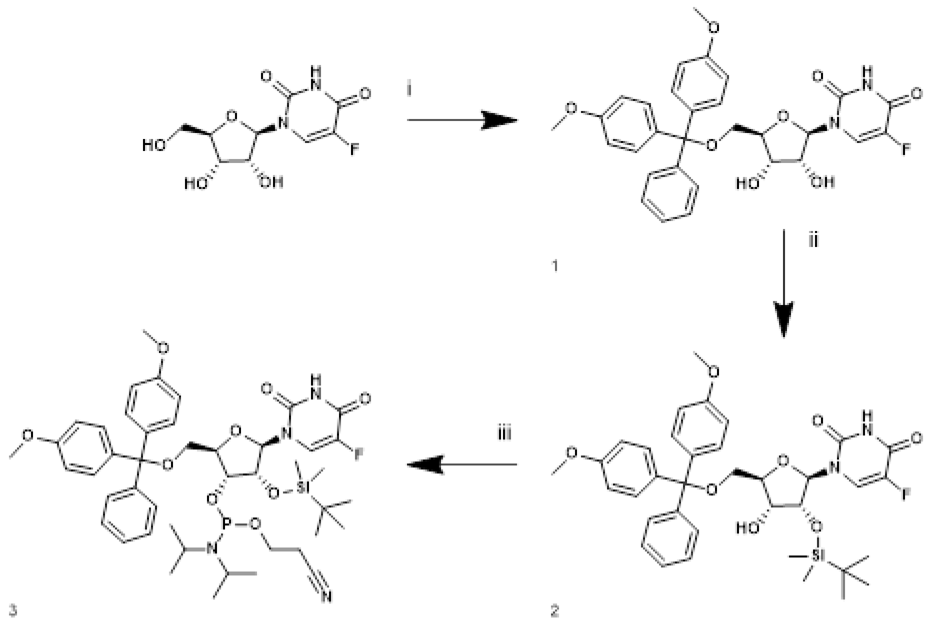

I) pyridine, 4,4'-Dimethoxytrityl chloride, r.t., 71%; II) $AgNO_3$ , pyridine, tert-Butyldimethylsilyl chloride, THF, r.t., 81%; III) DIPEA, THF, 2-Cyanoethyl N,N-diisopropylchlorophosphoramidite, 0°C to r.t., o.n., 92%.

**1-((2R,3R,4S,5R)-5-((bis(4-methoxyphenyl)(phenyl)methoxy)methyl)-3,4-dihydroxytetrahydrofuran-2-yl)-5-fluoropyrimidine-2,4(1H,3H)-dione** (**1**). In a round-bottom flask under a nitrogen atmosphere, 5-Fluorouridine (2.40 g, 9.20 mmol, 1.0 equiv) was dissolved in dry pyridine (20 mL, 0.40 M). 4,4′-Dimethoxytrityl chloride (3.42 g, 10.1 mmol, 1.1 equiv) was added portionwise, and the reaction mixture was stirred at room temperature for 16 h. Reaction progress was monitored by UPLC–MS. Upon complete consumption of the starting material, the reaction mixture was transferred to a separatory funnel, diluted with ethyl acetate, and washed with saturated aqueous ammonium chloride. The organic layer was collected, dried over anhydrous sodium sulfate, filtered, and concentrated under reduced pressure. The crude product was purified by automated flash chromatography (Biotage) using dichloromethane/methanol as the eluent (0–10% MeOH gradient) to afford the title compound as a white solid (3.70 g, 6.54 mmol, 71%). **MS (ESI)** m/z calc [M-H]-: 563.19, found 563.1. **$^1$H NMR (400 MHz, $CDCl_3$_$_3$mm)** δ 7.91 (d, *J* = 5.8 Hz, 1H), 7.74 – 7.64 (m, 1H), 7.33 – 7.16 (m, 8H), 6.86 – 6.77 (m, 4H), 5.91 (s, 1H), 5.29 (s, 1H), 4.44 – 4.35 (m, 2H), 4.23 – 4.18 (m, 1H), 3.75 (s, 1H), 3.73 (s, 6H), 3.47 – 3.42 (m, 2H).

**1-((2R,3R,4R,5R)-5-((bis(4-methoxyphenyl)(phenyl)methoxy)methyl)-3-((tert-butyldimethylsilyl)oxy)-4-hydroxytetrahydrofuran-2-yl)-5-fluoropyrimidine-2,4(1H,3H)-dione** (**2**). In a round-bottom flask under an argon atmosphere, compound **1** (3.70 g, 6.54 mmol, 1.0 equiv) was dissolved in dry THF (13 mL, 0.50 M). Pyridine (2.3 mL, 7.19 mmol, 1.1 equiv) and silver nitrate (1.22 g, 7.19 mmol, 1.1 equiv) were added sequentially, and the reaction mixture was stirred for 20 min. *tert*-Butyldimethylsilyl chloride (1.08 g, 7.19 mmol, 1.1 equiv) was then added, and the mixture was stirred at room temperature in the dark for 16 h. Reaction progress was monitored by UPLC–MS. Upon complete consumption of the starting material, the reaction mixture was filtered through a pad of Celite® and concentrated under reduced pressure. The crude product was purified by automated flash chromatography (Biotage) on silica gel using a cyclohexane/ethyl acetate gradient (0–30% EtOAc) to afford the title compound as a white solid (2.00 g, 5.28 mmol, 81%). **MS (ESI)** m/z calc [M-H]-: 677.28, found 677.3. **$^{1}$H NMR (400 MHz, DMSO)** δ 7.86 (d, *J* = 6.7 Hz, 1H), 7.32 (d, *J* = 8.0 Hz, 2H), 7.27 – 7.11 (m, 7H), 6.82 (d, *J* = 8.8 Hz, 4H), 5.69 – 5.63 (m, 1H), 5.07 (d, *J* = 6.2 Hz, 1H), 4.19 (t, *J* = 4.5 Hz, 1H), 4.04 – 3.97 (m, 1H), 3.97 – 3.90 (m, 2H), 3.66 (s, 5H), 3.30 (d, *J* = 4.6 Hz, 1H), 3.13 (dd, *J* = 10.7, 2.4 Hz, 1H), 0.79 (s, 9H), -0.00 (d, *J* = 2.2 Hz, 6H).

**(2R,3R,4R,5R)-2-((bis(4-methoxyphenyl)(phenyl)methoxy)methyl)-4-((tert-butyldimethylsilyl)oxy)-5-(5-fluoro-2,4-dioxo-3,4-dihydropyrimidin-1(2H)-yl)tetrahydrofuran-3-yl (2-cyanoethyl) diisopropylphosphoramidite** (**3**). In a round-bottom flask under a nitrogen atmosphere, compound **3** (1.385 g, 2.05 mmol, 1.0 equiv) and *N,N*-diisopropylethylamine (2.14 mL, 12.28 mmol, 6.0 equiv) were dissolved in dry THF (26 mL, 0.04 M). The reaction mixture was cooled to 0 °C, and 2-cyanoethyl *N,N*-diisopropylchlorophosphoramidite (1.83 mL, 8.18 mmol, 4.0 equiv) was added portionwise. The reaction mixture was allowed to warm to room temperature and stirred for 16 h. Reaction progress was monitored by UPLC–MS. Upon complete consumption of the starting material, the reaction was quenched at 0 °C by the addition of methanol, transferred to a separatory funnel, and washed with water. The organic layer was dried over anhydrous sodium sulfate, filtered, and concentrated under reduced pressure. The crude product was purified by automated flash chromatography on silica gel using a hexane/ethyl acetate gradient (0–100% EtOAc) to afford the title compound as a white powder (1.65 g, 1.88 mmol, 92%). **MS (ESI)** m/z calc [M-H]-: 877.39, found 877.6. **$^{31}$P NMR (202 MHz, $CD_3CN$)** δ 149.52, 149.14.

**5F-rFU labelled sample synthesis.**

All 5F-rFU labelled RNAs were synthesized and purified as described in the method part
